# The calcium channel subunit α_2_δ-3 organizes synapses via a novel activity-dependent, autocrine BMP signaling pathway

**DOI:** 10.1101/640664

**Authors:** Kendall M. Hoover, Scott J. Gratz, Kelsey A. Herrmann, Nova Qi, Alexander Liu, Jahci J. Perry-Richardson, Pamela J. Vanderzalm, Kate M. O’Connor-Giles, Heather T. Broihier

**Affiliations:** Department of Neurosciences, Case Western Reserve University School of Medicine, Cleveland, Ohio 44106; Department of Neuroscience, Brown University, Providence, Rhode Island 02912; Department of Biology, John Carroll University, University Heights, Ohio 44118

## Abstract

Synapses are highly specialized for neurotransmitter signaling, yet activity-dependent growth factor release also plays critical roles at synapses. While efficient neurotransmitter signaling is known to rely on precise apposition of release sites and neurotransmitter receptors, molecular mechanisms enabling high-fidelity growth factor signaling within the synaptic microenvironment remain obscure. Here we show that the auxiliary calcium channel subunit α2δ-3 promotes the function of a novel activity-dependent autocrine BMP signaling pathway at the Drosophila NMJ. α2δ proteins have conserved synaptogenic activity, although how they execute this function has remained elusive. We find that α2δ-3 provides an extracellular scaffold for autocrine BMP signaling, suggesting a new mechanistic framework for understanding α2δ’s conserved role in synapse organization. We further establish a transcriptional requirement for activity-dependent, autocrine BMP signaling in determining synapse density, structure, and function. We propose that activity-dependent, autocrine signals provide neurons with continuous feedback on their activity state and are thus well poised to modulate synapse structure and function.

## Introduction

Synapses are asymmetric cellular junctions underlying directional information transfer in the nervous system. Activity-dependent pathways are often invoked as regulators of synapse development, maintenance, plasticity, and elimination. While roles for activity-dependent cues at synapses are indisputable, the molecular links between activity and synapse differentiation remain poorly defined. To establish these connections, it is imperative to define when and where activity-dependent cues are released, and then how these extracellular signals impinge on synapse morphology and function.

Classical morphogens and neurotrophins have emerged as central regulators of synaptic development. A diverse collection of secreted cues including BMPs, Wnts, IGFs, FGFs, BDNF, Netrins, and Ephrins promote differentiation of the pre- or postsynaptic compartment (Cao et al., 2011; Glasgow et al., 2018; Johnson-Venkatesh and Umemori, 2010; Packard et al., 2002; Shen and Scheiffele, 2010; Terauchi et al., 2016). Such cues are often proposed to coordinate pre- and postsynaptic development by signaling across the synapse. Autocrine signals also serve important functions at the synapse by providing ongoing feedback to individual pre- or postsynaptic cells regarding their activity state (Herrmann and Broihier, 2018). As increasing numbers of synaptic signals with distinct functions and cellular origins are identified, the question arises of how neurons discriminate among these information channels. It is reasonable to hypothesize that proteins in the synaptic cleft serve crucial roles controlling the spatiotemporal availability of secreted proteins at the synapse to facilitate signal segregation.

α_2_δ proteins are well-positioned to impact the activity of synaptic signals. They are best studied as accessory calcium channel subunits and are composed of two disulfide-linked peptides: an entirely extracellular α_2_ peptide and a δ peptide predicted to be membrane-associated via a GPI link (Davies et al., 2010; Dolphin, 2018). They are important for voltage-gated calcium (Ca_V_) channel trafficking to synaptic terminals as well as for biophysical properties of Ca_V_ channels (Dolphin, 2018; Ferron et al., 2018; Kadurin et al., 2016).

Underscoring α_2_δ functions at mature synapses, α_2_δ proteins are cellular targets of Gabapentin (Field et al., 2006), a commonly prescribed treatment for epilepsy and neuropathic pain. All α_2_δ proteins contain a von Willebrand factor A (VWA) domain, a protein-protein interaction domain characteristic of ECM proteins such as matrilins and collagens (Whittaker and Hynes, 2002). Interestingly, α_2_δ proteins interact with synaptogenic proteins including Thrombo-spondin-1 (TSP-1) (Eroglu et al., 2009; Risher et al., 2018), suggesting functions beyond Ca_V_ channel localization and function. In line with this hypothesis, Drosophila α_2_δ-3, also called Straightjacket (Stj), is required for bouton morphogenesis at the embryonic neuromuscular junction (NMJ)—a function independent of the Ca_V_ channel Cacophany (Cac) (Kurshan et al., 2009). Functions for Drosophila α_2_δ-3 in subsequent aspects of NMJ differentiation are incompletely understood, but are reported to include synaptic assembly, synaptic function, and presynaptic homeostatic plasticity (Dickman et al., 2008; Ly et al., 2008; Wang et al., 2016).

In this study, we report a functional requirement for Drosophila α_2_δ-3 in activity-dependent autocrine BMP signaling. At the Drosophila NMJ, postsynaptic Gbb (Glass bottom boat), a BMP family member, is an early and permissive retrograde cue that initiates scaling growth of the NMJ (Berke et al., 2013; McCabe et al., 2003). We previously demonstrated that presynaptic Gbb is also trafficked to presynaptic terminals, where it is released in response to activity (James and Broihier, 2011; James et al., 2014). What is the function of this activity-dependent cue? Here we show that it is an autocrine signal required to maintain synapse structure and function. In the absence of presynaptic Gbb, active zone architecture, synaptic vesicle distribution, and baseline neurotransmitter release are all disrupted. Loss of the BMP-responsive transcription factor Mad results in similar phenotypes, arguing for a canonical BMP pathway. Embryonic bouton morphogenesis is also disrupted in BMP mutants, a phenotype identical to that seen in α_2_δ-3 mutants (Kurshan et al., 2009), prompting us to test for a link between BMP signaling and α_2_δ-3. Indeed, we find that structural and behavioral defects in α_2_δ-3 mutants are suppressed by activation of autocrine BMP signaling, suggesting that α_2_δ-3 mutant NMJs have reduced autocrine BMP signaling. Providing mechanistic insight into the function of α_2_δ-3 in autocrine BMP signaling, the extracellular α_2_ peptide of α_2_δ-3 prevents diffusion of Gbb following its activity-dependent release. We therefore propose that α_2_δ-3 is a key component of the microenvironment in the synaptic cleft that acts as a diffusion barrier to extracellular Gbb.

## Results

### A presynaptic, autocrine BMP pathway at the Drosophila NMJ

Seminal studies demonstrate that BMP signaling orchestrates NMJ development and physiology. Constitutive loss of BMP signaling causes a reduction in overall NMJ size as judged by bouton number—as well as ultrastructural defects, reduced evoked glutamate release, and impaired homeostatic plasticity (Goold and Davis, 2007; Marques et al., 2002; McCabe et al., 2003; 2004). These widespread defects raise the question of whether the phenotypes have a common root or, if instead, they reflect separable cell-specific roles for BMP signaling. Early work suggested at least partially separable pre- and postsynaptic BMP pathways; while expression of Gbb in the postsynaptic muscle rescues bouton number in *gbb* constitutive nulls, it does not rescue evoked neurotransmitter release. Expression of Gbb in the presynaptic neuron is required to restore proper glutamate release to *gbb* nulls (Goold and Davis, 2007; James et al., 2014; McCabe et al., 2003). These findings suggest that Gbb is released by the presynaptic motor neuron. Lending support to this idea, we found that Gbb is trafficked to presynaptic terminals, where it is subject to activity-dependent release (James et al., 2014).

We hypothesized that this presynaptic pool of Gbb regulates synapse formation or maintenance. To explore this idea, we first established the effect of constitutive loss of Gbb. As expected, overall NMJ size, as assessed by bouton number, is significantly decreased in *gbb* nulls **(Figure S1 A-B, H)**. Each bouton contains many synapses, or individual presynaptic glutamate release sites precisely aligned to postsynaptic glutamate receptor clusters. We utilized the ELKS-related protein Bruchpilot (Brp) as a presynaptic marker, GluRIII as a postsynaptic marker—and defined a synapse as a pair of Brp/GluRIII puncta (Fouquet et al., 2009; Kittel et al., 2006; Marrus et al., 2004; Wagh et al., 2006). We scored Brp-positive synapse density (synapse number/µm^2^) to exclude differences in synapse number arising as a secondary consequence of altered overall NMJ size. We find that loss of Gbb results in a 30% decrease in Brp-positive synapse density **(Figure 1 A-B, H)**, indicating that Gbb regulates synapse development.

**Figure 1.**
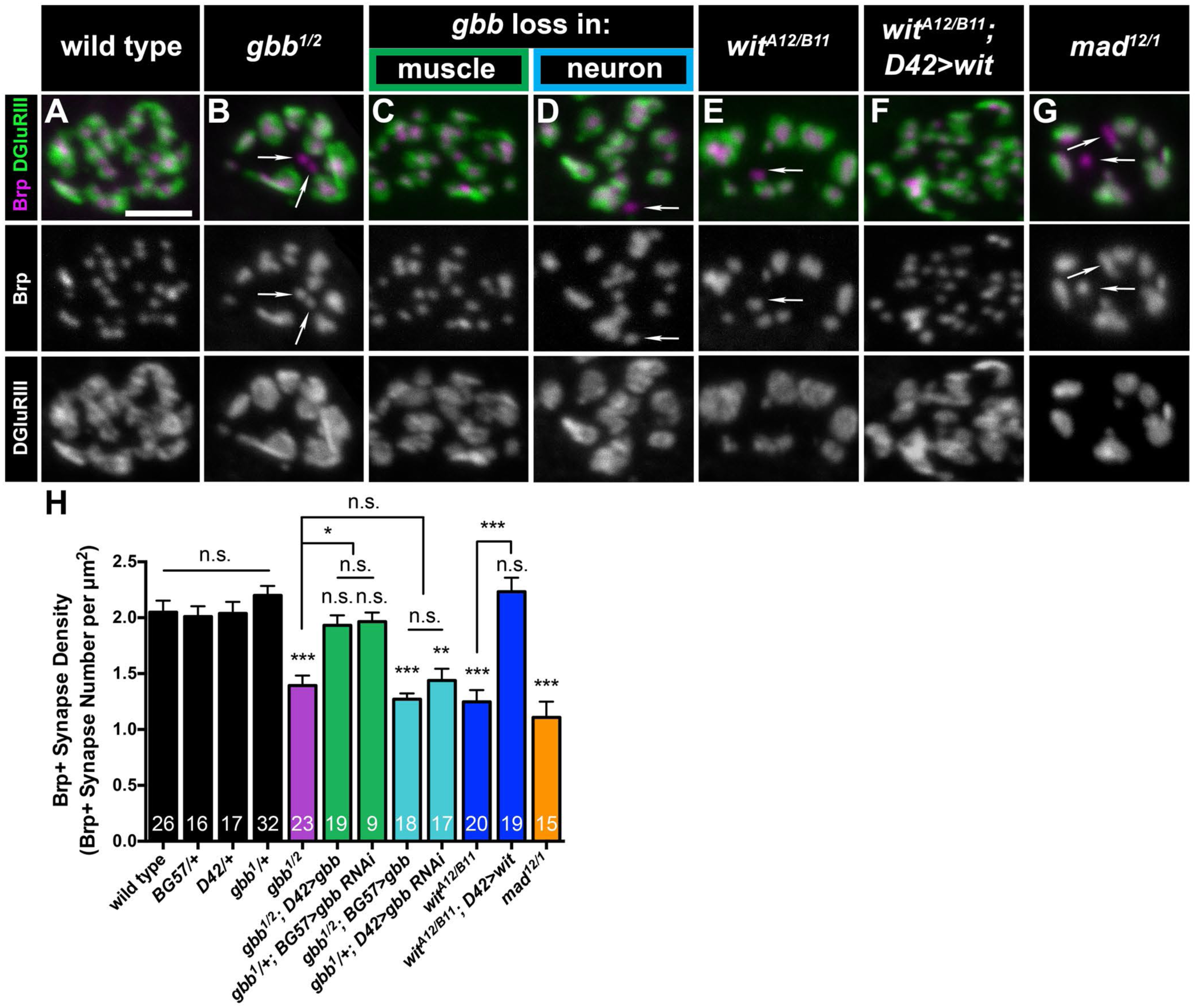
A presynaptic, autocrine BMP pathway at the Drosophila NMJ. (A-G) Representative z-projections of boutons of indicated genotypes labeled with Brp (magenta) and DGluRIII (green). Arrows denote unapposed Brp+ puncta. Individual channels are shown in grayscale below or next to merged images for clarity throughout all figures. Scale bar: 2 µm. (H) Quantification of Brp+ synapse density (number of Brp+ synapses per terminal bouton area in square microns). For all Brp+ synapse density analyses, n is the number of boutons scored. Wild type (*OregonR*): 2.1 ± 0.1, *BG57/+*: 2.0 ± 0.1, *D42/+*: 2.0 ± 0.1, *gbb^1^/+*: 2.2 ± 0.1, *gbb^1/2^*: 1.4 ± 0.1, *gbb^1/2^; D42>gbb*: 1.9 ± 0.1, *gbb^1^/+; BG57>gbb RNAi*: 2.0 ± 0.1, *gbb^1/2^; BG57>gbb*: 1.3 ± 0.1, *gbb^1^/+; D42>gbb RNAi*: 1.4 ± 0.1, *wit^A12/B11^*: 1.2 ± 0.1, *wit^A12/B11^; D42>wit*: 2.2 ± 0.1, *mad^12/1^*: 1.1 ± 0.1. Error bars are mean ± SEM. n.s., not significantly different. *, p<0.05; **, p<0.01; ***, p<0.001.

We defined the relevant cellular source of Gbb by selectively removing either pre- or postsynaptic ligand. To test necessity of the postsynaptic pool we used muscle-specific *gbb* RNAi (*gbb^1^/+; BG57>gbb RNAi*). In both this RNAi experiment and those described below, knock-down was performed in *gbb* heterozygotes. *gbb* heterozygosity does not impact synapse density in an otherwise wild-type background **(Figure 1 H)**. To confirm this RNAi-based approach and to test sufficiency of the presynaptic pool, we overexpressed Gbb in motor neurons in *gbb* nulls (*gbb^1/2^; D42>Gbb)*. As expected, muscle-derived ligand is required for NMJ expansion, since bouton number is significantly reduced in both backgrounds (Marques et al., 2002; McCabe et al., 2003) **(Figure S1 A, C, H)**. In contrast, we do not detect a change in Brp/GluRIII density **(Figure 1 A, C, H)**, indicating that muscle-derived Gbb does not otherwise regulate synapse development. We gained genetic access to motor neuron-derived Gbb using analogous strategies. We utilized motor neuron-specific RNAi (*gbb^1^/+; D42>gbb RNAi*) to probe a requirement for presynaptic ligand, while to test sufficiency of the postsynaptic ligand, we expressed Gbb in muscle in a *gbb* null (*gbb^1/2^; BG57>Gbb*). As expected, bouton number is unchanged in these backgrounds **(Figure S1 A, D, H)**, indicating that motor neuron-derived ligand does not regulate NMJ growth. However, the density of Brp/GluRIII puncta in both backgrounds is decreased comparably to *gbb* nulls **(Figure 1 A, D, H)**. Thus, the synapse density phenotype observed in *gbb* nulls is attributable to neuron-derived ligand.

Wishful thinking (Wit) is the neuronal Type II BMP receptor mediating Gbb function in NMJ growth (McCabe et al., 2004; Marques et al., 2003). We were curious if Wit is also required for Gbb’s synapse density function. As expected, loss of Wit results in fewer boutons **(Figure S1 A, E, H)**. In addition, we observe a 40% decrease in Brp-positive synapse density **(Figure 1 A, E, H)**, indicating that Wit regulates synapse development. To determine whether presynaptic Wit carries out this function, we assayed whether Wit expression in motor neurons in *wit* nulls (*wit^A12/B11^; D42>Wit*) rescues the decrease in synapse density. Indeed, synapse density is fully rescued by motor neuron expression of Wit **(Figure 1 A, F, H)**, indicating a presynaptic requirement. The genetic requirement of ligand and receptor in the presynaptic neuron, and not in the muscle, argues for an autocrine BMP signaling loop. Lastly, we were interested in whether this autocrine pathway requires Mad, the canonical transcription factor in the BMP pathway. Loss of Mad results in a 45% reduction in synapse density, supporting a canonical BMP pathway **(Figure 1 A, G-H)**. Together, these findings uncover a canonical, presynaptic and autocrine BMP pathway that promotes Brp-positive synapse density at the NMJ.

### Presynaptic, autocrine BMP signaling regulates active zone architecture and composition

In the preceding analysis, we had the impression that loss of autocrine BMP signaling results in larger and irregular Brp puncta **(Figure 1)**. When labeled with a C-terminal Brp antibody (nc82) and imaged in planar orientation at higher resolution, Brp puncta resolve into roughly 200 nm diameter rings **(Figure 2 A)** (Fouquet et al., 2009). Loss of key active zone cytomatrix components alter Brp ring geometry (Bruckner et al., 2017; Liu et al., 2011; Müller et al., 2012; Owald et al., 2010; 2012) making it a sensitive probe for assembly of the highly ordered cytomatrix. Thus, we analyzed Brp ring size and distribution utilizing extended resolution microscopy to gain insight into *gbb* mutant phenotypes.

**Figure 2.**
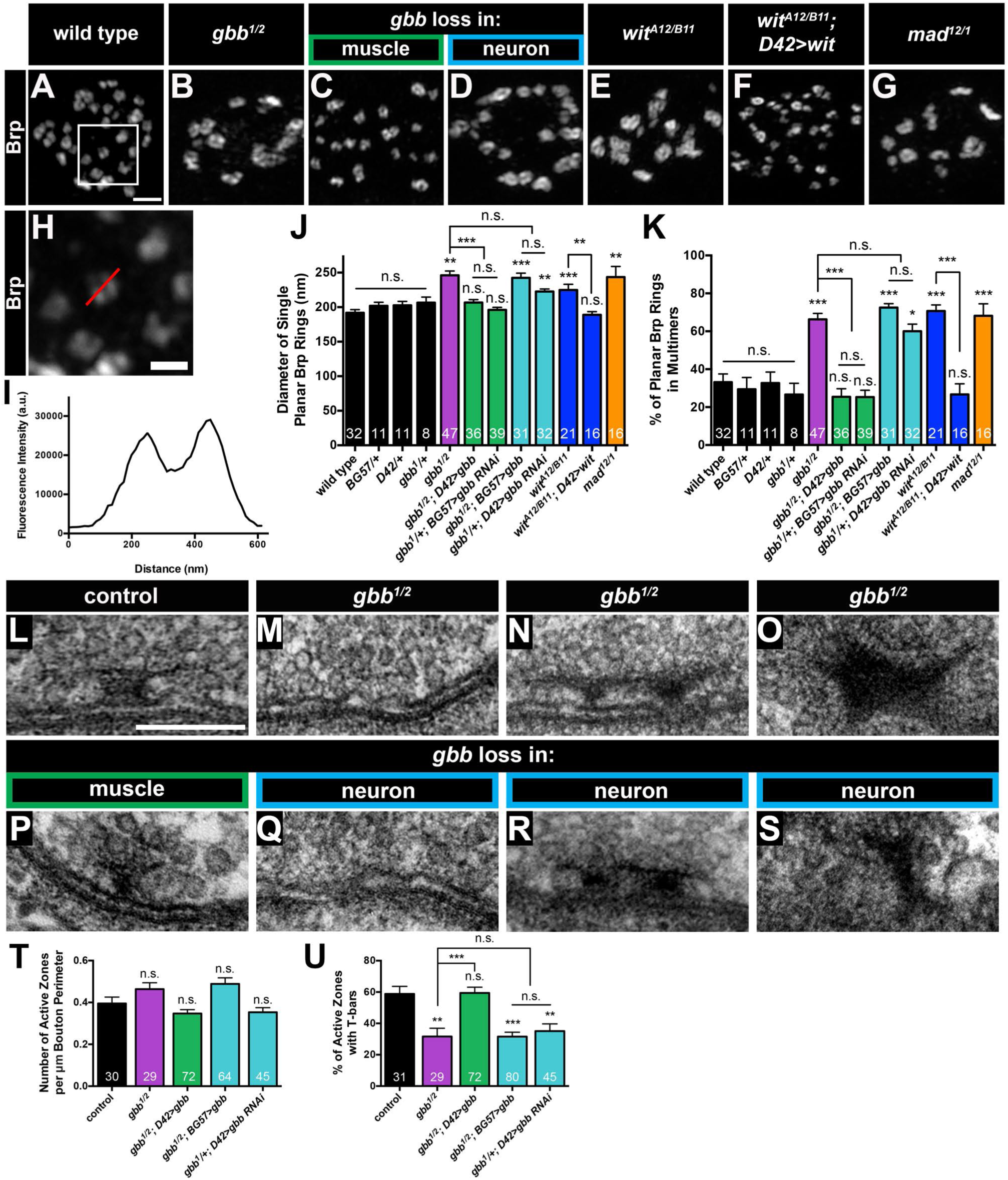
Presynaptic, autocrine BMP signaling regulates active zone architecture and composition. (A-G) Representative deconvolved z-projections of boutons of indicated genotypes labeled with anti-Brp. Scale bar: 1 µm. (H) Zoomed in view of the box in (A), demonstrating how Brp ring diameter was measured. The red line is typical of a manually drawn line through a single planar Brp ring. Scale bar: 200 nm. (I) Fluorescence intensity along the line in (H) was plotted, and Brp ring diameter was calculated as the distance between the two intensity maxima. (J) Quantification of single planar Brp ring diameters. For all Brp ring analyses, n is the number of boutons scored. Wild type (*OregonR*): 191.8 ± 4.7, *BG57/+*: 201.9 ± 5.2, *D42/+*: 202.6 ± 5.7, *gbb^1^/+*: 206.5 ± 8.8, *gbb^1/2^*: 246.1 ± 6.1, *gbb^1/2^; D42>gbb*: 206.8 ± 4.1, *gbb^1^/+; BG57>gbb RNAi*: 196.1 ± 3.5, *gbb^1/2^; BG57>gbb*: 242.4 ± 6.9, *gbb^1^/+; D42>gbb RNAi*: 222.6 ± 3.8, *wit^A12/B11^*: 224.8 ± 8.3, *wit^A12/B11^; D42>wit*: 189.0 ± 4.5, *mad^12/1^*: 243.6 ± 15.3. (K) Quantification of the percent of planar Brp rings within multimers, or groups of interconnected Brp rings. Wild type (*OregonR*): 33.2 ± 4.3, *BG57/+*: 29.4 ± 6.2, *D42/+*: 32.7 ± 5.8, *gbb^1^/+*: 26.7 ± 5.9, *gbb^1/2^*: 66.3 ± 3.2, *gbb^1/2^; D42>gbb*: 25.5 ± 4.2, *gbb^1^/+; BG57>gbb RNAi*: 25.3 ± 3.6, *gbb^1/2^; BG57>gbb*: 72.5 ± 2.1, *gbb^1^/+; D42>gbb RNAi*: 60.1 ± 3.7, *wit^A12/B11^*: 70.7 ± 3.2, *wit^A12/B11^; D42>wit*: 26.7 ± 5.6, *mad^12/1^*: 68.2 ± 6.3. (L-S) Representative transmission electron micrographs of active zones of the indicated genotypes. n is the number of active zones scored. Scale bar: 200 nm. (T) Quantification of the number of active zones per micron bouton perimeter. Control (*BG57/+*): 0.4 ± 0.0, *gbb^1/2^*: 0.5 ± 0.0, *gbb^1/2^; D42>gbb*: 0.3 ± 0.0, *gbb^1/2^; BG57>gbb*: 0.5 ± 0.0, *gbb^1^/+; D42>gbb RNAi*: 0.4 ± 0.0. (U) Quantification of the percent of active zones with T-bars. Control (*BG57/+*): 58.8 ± 4.8, *gbb^1/2^*: 31.6 ± 5.3, *gbb^1/2^; D42>gbb*: 59.4 ± 3.7, *gbb^1/2^; BG57>gbb*: 31.6 ± 2.8, *gbb^1^/+; D42>gbb RNAi*: 35.1 ± 4.6. Error bars are mean ± SEM. n.s., not significantly different. *, p<0.05; **, p<0.01; ***, p<0.001.

In controls, we find that the average diameter of isolated Brp rings is roughly 200 nm, consistent with published reports **(Figure 2 A, H, I, J)** (Bruckner et al., 2017; Kittel et al., 2006; Müller et al., 2012; Owald et al., 2010; 2012). In contrast, mean Brp ring diameter increases by 23% in *gbb* nulls **(Figure 2 B, J)**. We investigated the relevant cellular source of Gbb using cell-type specific knockdown and rescue genotypes described in detail above. We find that loss of neuronal Gbb results in increased Brp ring diameter analogous to that seen in *gbb* nulls, while loss of muscle-derived Gbb has no effect **(Figure 2 A-D, J)**. We next analyzed whether Wit and Mad also regulate cytomatrix architecture and find that loss of either one results in increased Brp ring diameter **(Figure 2 A, E, G, J)**. The *wit* mutant phenotype is rescued by neuronal expression of Wit, again pointing to an autocrine signaling pathway **(Figure 2 A, F, J)**. The increased Brp ring diameter in autocrine BMP mutants indicates that the pathway ensures precise construction and/or maintenance of the active zone cytomatrix.

Our analysis also revealed a clear defect in the distribution of Brp rings in autocrine BMP pathway mutants. In controls, the majority of Brp rings are separated from each other and observed as isolated rings, while in autocrine BMP mutants, many Brp rings are present in interconnected, disorganized clusters. We quantified this phenotype and found that roughly 31% of Brp rings are interconnected in controls, relative to 66% in *gbb* nulls **(Figure 2 A-B, K)**. This phenotype is apparent when presynaptic, but not postsynaptic, ligand is lost **(Figure 2 A, C-D, K)**, and likewise requires Mad and neuronal Wit **(Figure 2 A, E-G, K)**. These analyses suggest that autocrine BMP signaling regulates Brp allocation among active zones.

Together, these light-level analyses suggest that the observed 30% decrease in synapse density in autocrine BMP mutants may be explained by altered Brp distribution among active zones. In other words, some active zones may have excess Brp while others have none. Brp is an essential component of electron-dense T-bars visible at the EM level (Fouquet et al., 2009). Thus, if autocrine BMP signaling regulates Brp allocation to individual active zones, mutants might exhibit altered T-bar distribution. To test this hypothesis, we analyzed presynaptic ultrastructure in *gbb* mutants. We first asked if Gbb regulates the number of synapses, defined as electron-dense tightly apposed pre- and postsynaptic membranes. We scored active zone number per µm bouton perimeter in control, *gbb* null, as well as in animals with cell-type specific loss of Gbb and do not detect any alterations **(Figure 2 T)**. Thus, Gbb is not required to establish active zones. However, presynaptic Gbb promotes adhesion of pre- and postsynaptic synaptic membranes, since membrane ruffling increases in mutant backgrounds (from <5% in controls to roughly 20% in presynaptic Gbb mutants), consistent with membrane ruffling reported in constitutive BMP mutants (Aberle et al., 2003; McCabe et al., 2003).

In contrast, we find a marked reduction in the percentage of active zones containing T-bars. In our electron micrographs, 59% of control active zones have visible T-bars, in line with published work **(Figure 2 L, U)** (Graf et al., 2009). In contrast, T-bars are present at only 32% of *gbb* null active zones **(Figure 2 L, M, U)**. This phenotype tracks specifically with loss of the presynaptic ligand pool **(Figure 2 L, P, Q, U)**. Thus, presynaptic Gbb promotes T-bar localization to active zones. We observed two additional phenotypes in presynaptic Gbb mutants. First, we find an increase in active zones with multiple T-bars in *gbb* null and presynaptic *gbb* mutants **(Figure 2 N, R)**. And second, we observe membrane-detached aggregates of electron dense material **(Figure 2 O, S)**, which were never observed in controls, but have been reported with constitutive loss of the BMP pathway (Aberle et al., 2002; McCabe et al., 2003). We propose that these detached aggregates of T-bar material underlie the unopposed Brp puncta observed at the light level **(arrows in Figure 1 B, D, E, G)**. Taken together, these analyses indicate that autocrine BMP signaling regulates the architecture and composition of the presynaptic compartment.

### Presynaptic, autocrine BMP signaling regulates synaptic vesicle distribution

The preceding analyses demonstrate that autocrine BMP signaling regulates organization of the active zone cytomatrix. Might this pathway regulate additional features of the presynaptic compartment? We turned to the question of small synaptic vesicle (SSV) localization since tight regulation of vesicle number and distribution underpins presynaptic function. We analyzed SSV distribution using the vesicular glutamate transporter (VGLUT) as an SSV marker (Daniels et al., 2004). SSVs cluster at active zones for activity-dependent release and are enriched at presynaptic membranes, leaving bouton interiors relatively less populated with SSVs **(Figure 3 A)**. In contrast, in *gbb* mutants, we observed that SSVs are more uniformly distributed **(Figure 3 B)**. To quantify this phenotype, we assessed relative SSV distribution by drawing a line through the bouton center and analyzing the fluorescence intensity profile along the line. We defined proper SSV distribution as corresponding to a 50% decrease in VGLUT fluorescence intensity in the bouton interior **(Figure 3 H, I)**. We analyzed only boutons of at least 2 μM diameter to avoid small boutons lacking well-defined SSV distributions. We also calculated integrated VGLUT intensity as an estimate of overall vesicle number. Indicating that BMP signaling does not regulate overall SSV number at the NMJ, integrated VGLUT fluorescence intensity is not altered in any BMP pathway backgrounds **(Figure 3 K)**.

**Figure 3.**
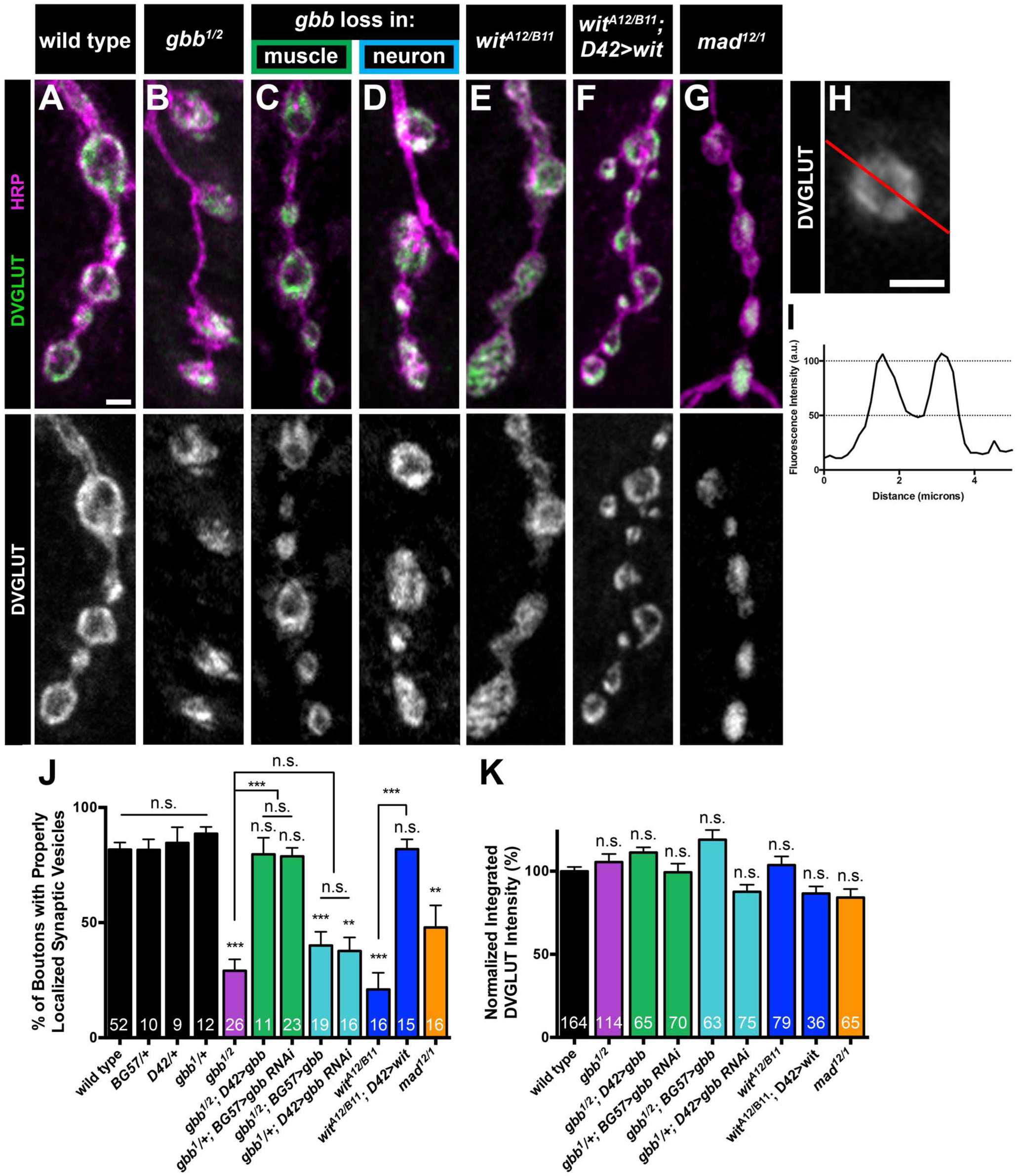
Presynaptic, autocrine BMP signaling regulates synaptic vesicle distribution. (A-G) Representative z-projections of boutons of the indicated genotypes labeled with DVGLUT (green) and HRP (magenta). Scale bar: 2 µm. (H) The red line represents a typical line drawn through a bouton larger than 2 µm in diameter. Scale bar: 2 µm. (I) The fluorescence intensity graph along the line (H) was created as shown. If the two intensity maxima were each greater than twice the minimum, the bouton had properly distributed synaptic vesicles. (J) Quantification of the percent of boutons properly distributed synaptic vesicles. For all DVGLUT localization analyses, n is the number of NMJs scored. Wild type (*OregonR*): 81.8 ± 3.1, *BG57/+*: 81.6 ± 4.6, *D42/+*: 84.6 ± 6.8, *gbb^1^/+*: 88.6 ± 3.0, *gbb^1/2^*: 29.1 ± 4.9, *gbb^1/2^; D42>gbb*: 79.7 ± 7.2, *gbb^1^/+; BG57>gbb RNAi*: 78.8 ± 3.7, *gbb^1/2^; BG57>gbb*: 40.1 ± 5.9, *gbb^1^/+; D42>gbb RNAi*: 37.7 ± 5.9, *wit^A12/B11^*: 20.9 ± 7.3, *wit^A12/B11^; D42>wit*: 81.9 ± 4.3, *mad^12/1^*: 47.9 ± 9.6. (K) Quantification of integrated DVGLUT intensity normalized to controls. For all integrated DVGLUT analyses, n is the number of boutons scored. Wild type (*OregonR*): 100.0 ± 2.6, *gbb^1/2^*: 105.5 ± 4.8, *gbb^1/2^; D42>gbb*: 111.3 ± 3.1, *gbb^1^/+; BG57>gbb RNAi*: 99.3 ± 5.2, *gbb^1/2^; BG57>gbb*: 118.9 ± 5.8, *gbb^1^/+; D42>gbb RNAi*: 87.6 ± 4.3, *wit^A12/B11^*: 103.7 ± 5.2, *wit^A12/B11^; D42>wit*: 86.6 ± 4.3, *mad^12/1^*: 84.2 ± 5.1. Error bars are mean ± SEM. n.s., not significantly different. **, p<0.01; ***, p<0.001.

In contrast, whereas 82% of control boutons have peripherally localized SSVs, only 29% of boutons in *gbb* nulls exhibit a peripheral SSV distribution **(Figure 3 A-B, J)**. Moreover, it is neuron-derived Gbb that regulates SSV distribution as genotypes with neuron-specific Gbb loss display aberrant SSV distribution indistinguishable from *gbb* nulls, while muscle-specific loss maintains peripheral enrichment of SSVs **(Figure 3 A, C-D, J)**. A similar requirement for neuronal Gbb in bouton-wide SSV distribution is also apparent in our EM analysis **(Figure S2)**. Thus, neuronal Gbb promotes normal SSV distribution. We next asked whether neuronal Wit and Mad are likewise involved. Supporting a function for Wit in regulating SSV distribution, only 21% of boutons in *wit* nulls have peripheral SSV distribution. This phenotype is rescued by neuronal expression of Wit, demonstrating a presynaptic requirement **(Figure 3 A, E-F, J)**. We find that loss of Mad results in a more intermediate phenotype, with 48% of boutons displaying normal SSV distribution **(Figure 3 A, G, J)**. Together, these findings argue that presynaptic, autocrine Gbb signaling regulates SSV distribution. Interestingly, LOF mutations in the conserved synaptic adhesion molecule Neurexin-1 display the same phenotype (Rui et al., 2017), suggesting that large-scale spatial organization of SSVs within boutons could be a common function of synapse organizing pathways.

### Presynaptic Gbb is necessary for baseline synaptic function

Genetic rescue experiments establish that neuronal Gbb is sufficient to restore normal baseline glutamate release to *gbb* nulls (Goold and Davis, 2007; James et al., 2014; McCabe et al., 2003). While these studies demonstrate that neuronal Gbb can support normal transmission, they do not demonstrate that neuronal Gbb normally does serve this function. The abnormalities in active zone structure and vesicle distribution that we observe with loss of neuronal Gbb are consistent with a role for this ligand pool in transmission.

To address this question directly, we measured spontaneous and evoked synaptic potentials via intracellular recordings in *gbb* RNAi backgrounds. We find that mutants with Gbb removed selectively from motor neurons exhibit a 53% reduction in evoked EJP amplitude, while mutants with Gbb removed from the muscle have a 26% reduction **(Figure 4 A-E)**. The 26% reduction in EJP amplitude with muscle-specific Gbb knockdown was expected as these NMJs display a 22% reduction in overall NMJ size without increased synapse density **(Figures 1 and S1)**. In contrast, the 53% reduction in EJP amplitude with loss of neuron-specific Gbb knockdown cannot be explained by a change in gross NMJ morphology since bouton number is unchanged in this background **(Figure S1 A, D, H)**. Instead, these findings support a specific requirement for presynaptic Gbb in synaptic function. Indeed, quantal content is significantly reduced upon presynaptic loss of Gbb **(Figure 4 G)**, indicating aberrant evoked glutamate release consistent with the observed structural deficits. These findings extend previous genetic rescue data (Goold and Davis, 2007; James et al., 2014; McCabe et al., 2003) to establish a functional requirement for presynaptic Gbb in evoked neurotransmitter release.

**Figure 4.**
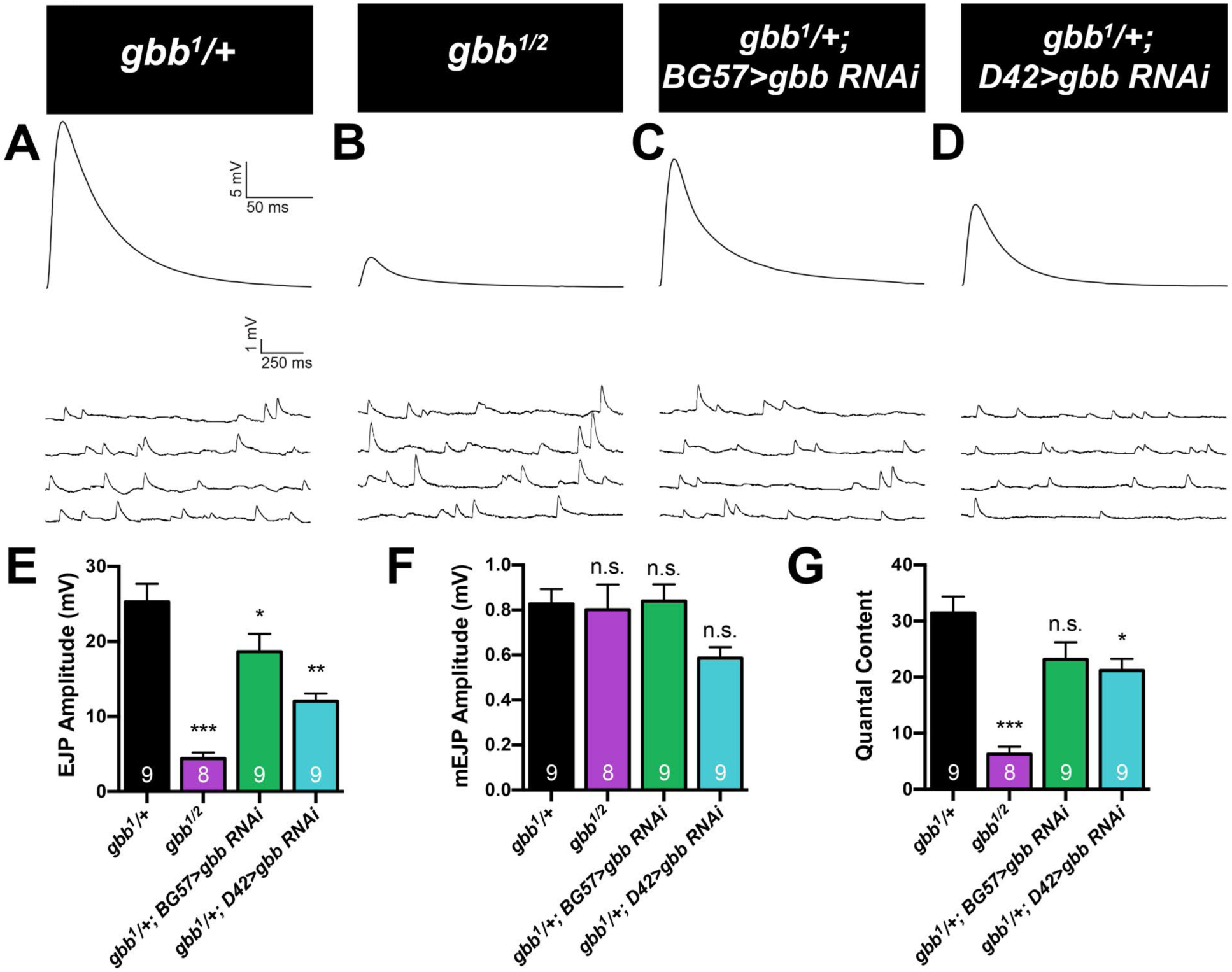
Presynaptic Gbb is necessary for baseline synaptic function. (A-D) Representative EJP (upper) and mEJP (lower) traces for the indicated genotypes. (E) Quantification of EJP amplitude. For all electrophysiological experiments, n is the number of cells recorded. *gbb^1^/+*: 25.3 ± 2.4, *gbb^1/2^*: 4.4 ± 0.8, *gbb^1^/+; BG57>gbb RNAi*: 18.6 ± 2.4, *gbb^1^/+; D42>gbb RNAi*: 12.0 ± 1.0. (F) Quantification of mEJP amplitude. *gbb^1^/+*: 0.8 ± 0.1, *gbb^1/2^*: 0.8 ± 0.1, *gbb^1^/+; BG57>gbb RNAi*: 0.8 ± 0.1, *gbb^1^/+; D42>gbb RNAi*: 0.6 ± 0.0. (G) Quantal content for each genotype was calculated by dividing average EJP amplitude by average mEJP amplitude. *gbb^1^/+*: 31.4 ± 2.9, *gbb^1/2^*: 6.3 ± 1.3, *gbb^1^/+; BG57>gbb RNAi*: 23.2 ± 3.1, *gbb^1^/+; D42>gbb RNAi*: 21.2 ± 2.1. Error bars are mean ± SEM. n.s., not significantly different. *, p<0.05; **, p<0.01; ***, p<0.001.

### Ongoing autocrine BMP signaling organizes the presynaptic compartment

Gbb and Mad are required exclusively during the first instar larval stage for NMJ growth (Berke et al., 2013), arguing for an early critical period for the retrograde pro-growth cue. To define the temporal requirement of presynaptic Gbb in organizing the presynaptic compartment, we utilized the GeneSwitch Gal4 system (GS-Gal4) for inducible expression of Gbb in all neurons via elav-GS-Gal4 (Osterwalder et al., 2001). Our aim was to define the critical period of autocrine Gbb signaling in active zone organization by inducing Gbb expression in neurons in *gbb* nulls at defined stages of larval development. Thus, we first tested if proper synapse density is rescued by inducing Gbb in neurons during the second instar larval stage (L2) and find that it is restored to wild-type levels **(Figure 5 A-B, D)**. Moreover, later induction of Gbb at third instar (L3) fully rescues the *gbb* null phenotype **(Figure 5 A, C, D)**. Supporting the temporal distinction between an early pro-growth signal and a later synapse-organizing signal, bouton number is not rescued with L2 or L3 Gbb induction **(Figure 5 E)**. We also measured planar Brp ring diameter in *gbb; Elav-GS>Gbb* animals and find that inducing Gbb at either L2 or L3 restores wild-type Brp ring diameter **(Figure 5 F-H, I)**. Finally, the interconnected Brp ring phenotype of *gbb* nulls is also completely rescued by L2 or L3 induction of Gbb **(Figure 5 F-H, J)**. These findings are consistent with prior work indicating a later larval requirement for Mad in neurotransmission (Berke et al., 2013) and further imply that inducing autocrine BMP signaling after synapses have formed is sufficient to correct aberrant synapse structure.

**Figure 5.**
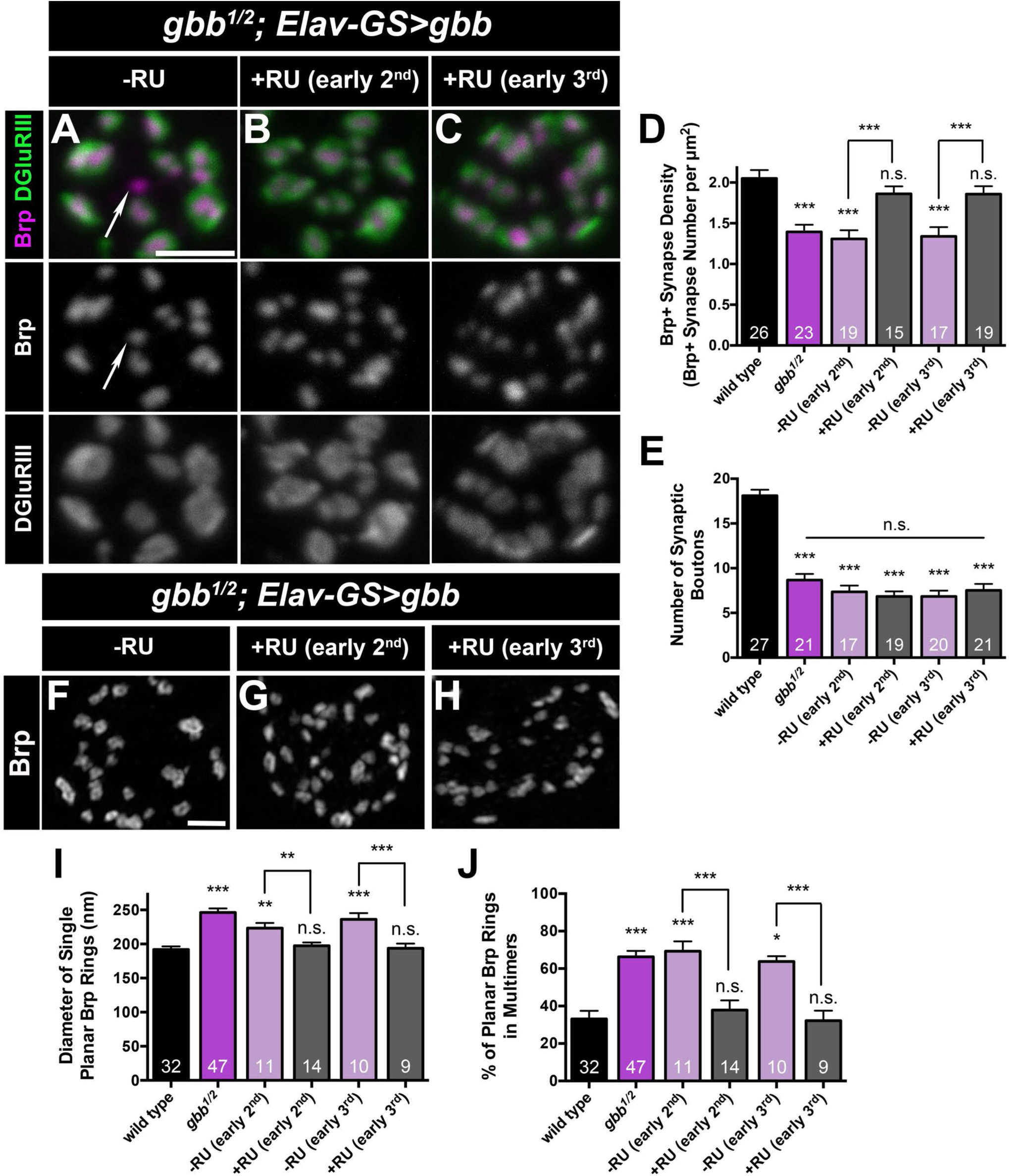
Ongoing autocrine BMP signaling organizes the presynaptic compartment. (A-C) Representative z-projections of boutons labeled with labeled with Brp (magenta) and DGluRIII (green) of *gbb^1/2^* mutants with transgenic *UAS-gbb* under the control of *ElavGeneSwitch*. Arrows denote unapposed Brp+ puncta. Scale bar: 2 µm. (D) Quantification of Brp+ synapse density. Wild type (*OregonR*): 2.1 ± 0.1, *gbb^1/2^*: 1.4 ± 0.1, -RU (early 2^nd^): 1.3 ± 0.1, +RU (early 2^nd^): 1.9 ± 0.1, -RU (early 3^rd^): 1.3 ± 0.1, +RU (early 3^rd^): 1.9 ± 0.1. (E) Quantification of the number of boutons. n is the number of NMJs scored. Wild type (*OregonR*): 18.1 ± 0.7, *gbb^1/2^*: 8.7 ± 0.7, -RU (early 2^nd^): 7.4 ± 0.7, +RU (early 2^nd^): 6.8 ± 0.6, -RU (early 3^rd^): 6.9 ± 0.6, +RU (early 3^rd^): 7.5 ± 0.7. (F-H) Representative z-projections of boutons of the indicated genotypes and treatments labeled with anti-Brp. Scale bar: 1 µm. (I) Quantification of single planar Brp ring diameters. Wild type (*OregonR*): 191.8 ± 4.7, *gbb^1/2^*: 246.1 ± 6.1, -RU (early 2^nd^): 223.3 ± 7.5, +RU (early 2^nd^): 197.4 ± 4.8, -RU (early 3^rd^): 236.2 ± 9.0, +RU (early 3^rd^): 193.8 ± 6.7. (J) Quantification of the percent of planar Brp rings in multimers. Wild type (*OregonR*): 33.2 ± 4.3, *gbb^1/2^*: 66.3 ± 3.2, -RU (early 2^nd^): 69.3 ± 5.3, +RU (early 2^nd^): 37.9 ± 5.2, -RU (early 3^rd^): 63.8 ± 2.9, +RU (early 3^rd^): 32.2 ± 5.3. Error bars are mean ± SEM. n.s., not significantly different. *, p<0.05; **, p<0.01; ***, p<0.001.

Together these findings argue that early retrograde Gbb signaling initiates scaling growth of the NMJ while continuous autocrine Gbb signaling organizes presynaptic terminals. The separable temporal requirements raise the possibility that the autocrine pathway is the predominant BMP pathway active at the L3 NMJ. In this event, autocrine signaling might more efficiently drive BMP signal transduction at this stage. We tested this hypothesis by comparing the effectiveness of pre- and postsynaptic ligand to drive nuclear pMad localization. We find that pMad localization in motor neuron nuclei is strongly reduced in *gbb* nulls **(Figure S3 A-B, E)**. And in accordance with our model, while expression of Gbb in the muscle weakly rescues pMad localization in *gbb* nulls, neuronal Gbb provides significantly greater rescue **(Figure S3 A-E)**. These results argue that autocrine Gbb signaling is the primary BMP pathway at the L3 stage.

### Loss of Gbb phenocopies loss of α_2_δ-3 at the embryonic NMJ

Our findings argue that Gbb regulates synapse maintenance; however, they do not address whether it regulates initial synapse formation. At 21 hours after egg laying (AEL), embryonic NMJs display hallmarks of mature NMJs such as rounded boutons and localized presynaptic components, including Brp (Rheuben et al., 1999; Yoshihara et al., 1997). Thus, to address whether Gbb is required for embryonic synapse formation, we assessed Brp localization in *gbb* mutants. In wild type, we find approximately 40 Brp-positive puncta at NMJ 6/7 at 21 h AEL **(Figure 6 A, E)**. We do not find a reduction in the number of Brp-positive puncta in either *gbb* or *wit* null embryos **(Figure 6 A-C, E)**, suggesting that Gbb signaling is not required for initial synapse formation. While these NMJs contain an appropriate number of Brp puncta, their gross morphology is markedly abnormal. Whereas wild-type NMJs at this stage are characterized by clusters of boutons, NMJs in *gbb* and *wit* mutants are elongated with fewer, smaller boutons. Indeed, we find a roughly 50% reduction in bouton number in *gbb* and *wit* mutants relative to controls **(Figure 6 F)**. The motor neuron-specific and muscle-specific drivers in our larval analysis did not enable us to map the cellular requirement of Gbb in this context; however, these findings demonstrate an overall function for BMP signaling in embryonic bouton formation.

**Figure 6.**
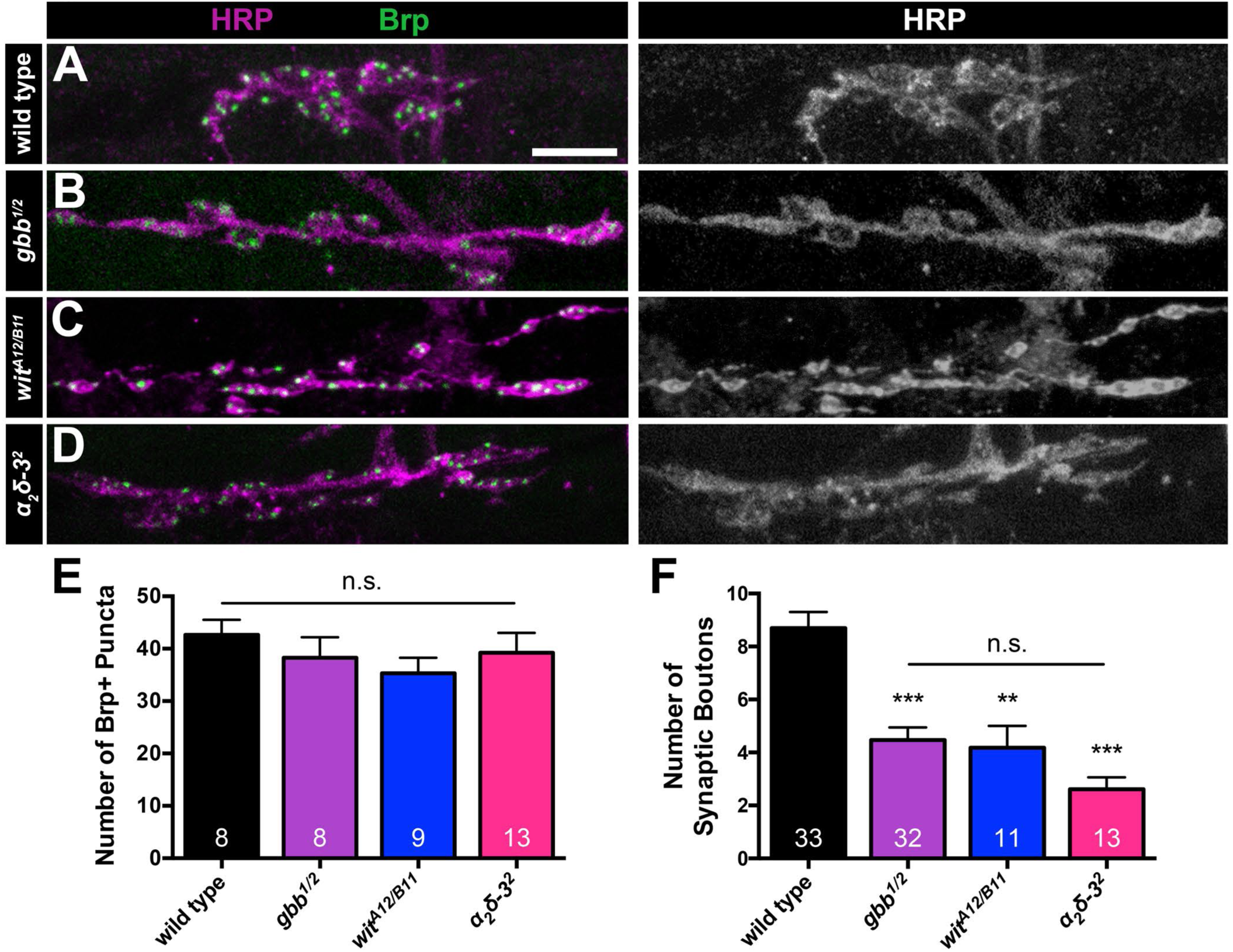
Loss of Gbb phenocopies loss of α_2_δ-3 at the embryonic NMJ. (A-D) Representative z-projections of embryonic NMJs of the indicated genotypes labeled with Brp (green) and HRP (magenta). Scale bar: 2 µm. (E) Quantification of the number of Brp+ puncta. For all embryonic experiments, n is number of NMJs scored. Wild type (*OregonR*): 42.6 ± 2.9, *gbb^1/2^*: 38.3 ± 3.9, *wit^A12/B11^*: 35.3 ± 3.0, *α_2_δ-3^2^*: 39.2 ± 3.8. (F) Quantification of the number of synaptic boutons. Wild type (*OregonR*): 8.7 ± 0.6, *gbb^1/2^*: 4.5 ± 0.5, *wit^A12/B11^*: 4.2 ± 0.8, *α_2_δ-3^2^*: 2.6 ± 0.4. Error bars are mean ± SEM. n.s., not significantly different. **, p<0.01; ***, p<0.001.

This embryonic phenotype caught our attention because it mirrors the phenotype displayed by strong *α_2_δ-3* alleles **(Figure 6 D-F)** (Kurshan et al., 2009). α_2_δ-3 is an auxiliary Ca^2+^ channel subunit that traffics and localizes Cacophany (Cac), the Drosophila α_1_ subunit of mammalian N- and P/Q-type voltage-gated calcium channels, to active zones (Dickman et al., 2008). α_2_δ-3 proteins contain two disulfide-linked peptides, α_2_ and δ. While the C-terminal δ peptide is membrane-associated and necessary for α_1_ subunit association, the large, heavily glycosylated N-terminal α_2_ domain is exclusively extracellular (Dolphin, 2018). Intriguingly, α_2_δ proteins are also conserved regulators of synapse formation and function (Dolphin, 2018; Eroglu et al., 2009; Hoppa et al., 2012; Pirone et al., 2014; Tong et al., 2017; Wang et al., 2016; 2017). Yet how α_2_δ proteins carry out their distinct functions in Ca^2+^ channel localization versus synapse formation is not well understood.

### Gbb and α_2_δ-3 have related functions in presynaptic organization

The phenotypic similarity between BMP mutants and *α_2_δ-3* mutants at embryonic NMJs raised the possibility of an underlying functional interaction. Hence, we tested if loss of α_2_δ-3 results in presynaptic deficits, making use of well-characterized *α_2_δ-3* alleles. *α_2_δ-3^DD106^* is a null allele; *α_2_δ-3^DD196^* is a hypomorphic allele harboring a stop codon following the α_2_ peptide; and *α_2_δ-3^k10814^* (here called *α_2_δ-3^k^*) is a hypomorphic P element allele with reduced expression of wild-type protein (Dickman et al., 2008; Kurshan et al., 2009) **(Figure 7 E)**. Importantly, while bouton formation is blocked in *α_2_δ-3^DD106^* homozygotes, *α_2_δ-3^DD196^* mutants have normal embryonic NMJs, suggesting a specific role for the extracellular α_2_ peptide in bouton morphogenesis (Kurshan et al., 2009). *α_2_δ-3^DD106^* homozygotes are late-stage embryonic lethal, so to characterize presynaptic organization at the L3 stage, we analyzed this allele in trans to the weak *α_2_δ-3^k^* allele, as well as *α_2_δ-3^DD196^* in trans to *α_2_δ-3^k^*. Consistent with α_2_δ-3’s role in Ca^2+^ channel localization, both allelic combinations *(α_2_δ-3^DD106/k^* and *α_2_δ-3^DD196/k^*) display strong reductions of Cac at active zones **(Figure S4 A-D)**.

**Figure 7.**
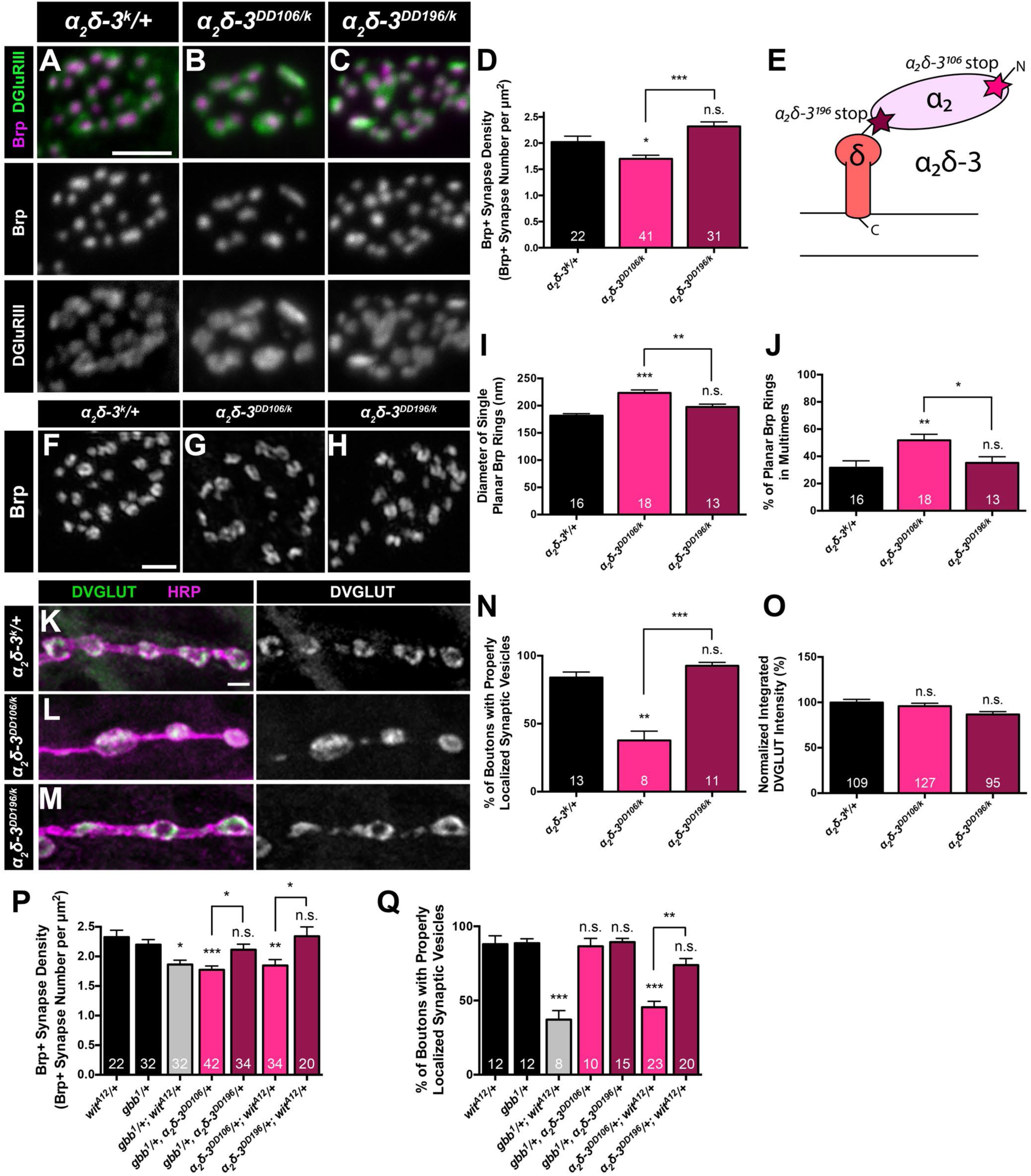
Gbb and α_2_δ-3 have related functions in presynaptic organization. (A-C) Representative z-projections of boutons of the indicated genotypes labeled with Brp (magenta) and DGluRIII (green). Scale bar: 2 µm. (D) Quantification of the Brp+ synapse density. *α_2_δ-3^k10814^/+*: 2.0 ± 0.1, *α_2_δ-3^DD106/k10814^*: 1.7 ± 0.1, *α_2_δ-3^DD196/k10814^*: 2.3 ± 0.1. (E) Schematic of *α_2_δ-3* subunit of a voltage-dependent Ca^2+^ channel. Stars mark the locations of the stop codons in *α_2_δ-3^DD106^* and *α_2_δ-3^DD196^* alleles. (F-H) Representative deconvolved z-projections of boutons of the indicated genotypes labeled with anti-Brp. Scale bar: 1 µm. (I) Quantification of single planar Brp ring diameters. *α_2_δ-3^k10814^/+*: 181.6 ± 3.8, *α_2_δ-3^DD106/k10814^*: 223.4 ± 5.5, *α_2_δ-3^DD196/k10814^*: 197.5 ± 5.4. (J) Quantification of the percent of planar Brp rings in multimers in boutons. *α_2_δ-3^k10814^/+*: 31.6 ± 5.1, *α_2_δ-3^DD106/k10814^*: 51.8 ± 4.4, *α_2_δ-3^DD196/k10814^*: 35.2 ± 4.5. (K-M) Representative z-projections of boutons of the indicated genotypes labeled with DVGLUT (green) and HRP (magenta). Scale bar: 2 µm. (N) Quantification of the percent of boutons exhibiting properly localized synaptic vesicles. *α_2_δ-3^k10814^/+*: 83.9 ± 4.1, *α_2_δ-3^DD106/k10814^*: 37.7 ± 6.9, *α_2_δ-3^DD196/k10814^*: 92.6 ± 2.5. (O) Quantification of integrated DVGLUT intensity compared to proper controls. *α_2_δ-3^k10814^/+*: 99.8 ± 3.5, *α_2_δ-3^DD106/k10814^*: 95.8 ± 3.2, *α_2_δ-3^DD196/k10814^*: 86.7 ± 3.1. (P) Quantification of Brp+ synapse density. *wit^A12^/+*: 2.3 ± 0.1, *gbb^1^/+*: 2.2 ± 0.1, *gbb^1^/+; wit^A12^/+*: 1.9 ± 0.1, *gbb^1^/+, α_2_δ-3^DD106^/+*: 1.8 ± 0.1, *gbb^1^/+, α_2_δ-3^DD196^/+*: 2.1 ± 0.1, *α_2_δ-3^DD106^/+; wit^A12^/+*: 1.8 ± 0.1, *α_2_δ-3^DD196^/+; wit^A12^/+*: 2.3 ± 0.2. (Q) Quantification of the percent of boutons exhibiting properly localized synaptic vesicles. *wit^A12^/+*: 88.0 ± 5.7, *gbb^1^/+*: 88.6 ± 3.0, *gbb^1^/+; wit^A12^/+*: 37.1 ± 6.1, *gbb^1^/+, α_2_δ-3^DD106^/+*: 86.6 ± 5.3, *gbb^1^/+, α_2_δ-3^DD196^/+*: 89.4 ± 2.5, *α_2_δ-3^DD106^/+; wit^A12^/+*: 45.4 ± 4.0, *α_2_δ-3^DD196^/+; wit^A12^/+*: 73.9 ± 4.4. Error bars are mean ± SEM. n.s., not significantly different. *, p<0.05; **, p<0.01; ***, p<0.001.

We next assessed presynaptic organization in *α_2_δ-3^DD106/k^* and *α_2_δ-3^DD196/k^* animals. We find that *α_2_δ-3^DD106/k^* mutant NMJs have synaptic defects analogous to those observed in presynaptic Gbb mutants. Specifically, the density of Brp/GluRIII pairs decreases **(Figure 7 A-D)**, while the diameter of isolated planar Brp rings and the percentage of interconnected Brp rings increases **(Figure 7 F-J)**. The observed decrease in synapse density in *α_2_δ-3^DD106/k^* mutants is similar to that reported for another strong *α_2_δ-3* allelic combination (Dickman et al., 2008). *α_2_δ-3^DD106/k^* mutants also display altered SSV distribution akin to presynaptic Gbb mutants **(Figure 7 K-O)**. In contrast, presynaptic organization appears normal in *α_2_δ-3^DD196/k^* mutants **(Figure 7 A-O)**. These findings argue that α_2_δ-3 regulates active zone cytomatrix architecture and SSV distribution and further suggest a selective requirement for the extracellular α_2_ domain.

Given the phenotypic similarities, we assayed for dominant genetic interactions between BMP pathway mutants and α_2_δ-3 mutants. In line with a common pathway, *α_2_δ-3^DD106^, gbb^1^* double heterozygotes and *α_2_δ-3^DD106^*, *wit^A12^* double heterozygotes display decreased Brp-positive synapse density similar to individual single mutants **(Figure 7 P)**. Notably, the strength of the interaction between *α_2_δ-3^DD106^* and *gbb* or *wit* is as strong as the genetic interaction between *gbb* and *wit* **(Figure 7 P)**. Moreover, *α_2_δ-3^DD106^*, *wit^A12^* double heterozygotes have deficits in SSV distribution akin to that of the single mutants **(Figure 7 Q)**. Surprisingly, *α_2_δ-3^DD106^, gbb^1^* double heterozygotes have normal SSV distribution **(Figure 7 Q)**, hinting at genetic redundancy. Together, these genetic interactions suggest that α_2_δ-3 functions with the autocrine BMP signaling pathway.

### Gbb overexpression in motor neurons suppresses *α_2_δ-3* mutant phenotypes

α_2_δ proteins are large, glycosylated proteins residing in the synaptic cleft (Dolphin, 2018), raising the possibility that α_2_δ-3 influences BMP signaling at the extracellular level. Moreover, our phenotypic analysis of α_2_δ-3 alleles supports a specific requirement for the extracellular α_2_ peptide in autocrine BMP signaling. We considered two hypotheses for how α_2_δ-3 and BMP signaling might interact. First, α_2_δ-3 might be required for autocrine BMP signaling, perhaps by serving as an obligate co-receptor. Alternatively, α_2_δ-3 might act in the extracellular space to potentiate autocrine BMP signaling by regulating Gbb levels or diffusion.

To distinguish between these hypotheses, we tested if over-activation of autocrine BMP signaling is sufficient to suppress *α_2_δ-3* phenotypes. We reasoned that if α_2_δ-3 facilitates autocrine BMP pathway activity by promoting its extracellular availability, then elevated Gbb release might suppress *α_2_δ-3* mutant phenotypes. In contrast, if α_2_δ-3 is an essential component of the BMP pathway, then elevated Gbb release is not predicted to ameliorate *α_2_δ-3* mutant phenotypes. We find that the decrease in Brp-positive synapse density observed in *α_2_δ-3^DD106/k^* animals is suppressed by overexpression of Gbb in motor neurons **(Figure 8 A-D)**. Similarly, proper SSV enrichment around the bouton perimeter is restored to *α_2_δ-3* mutants by Gbb overexpression in motor neurons **(Figure 8 E-H)**, suggesting that this phenotype is caused by reduced autocrine Gbb signaling. Together, these findings are consistent with the hypothesis that α_2_δ-3 normally facilitates autocrine BMP signaling.

**Figure 8.**
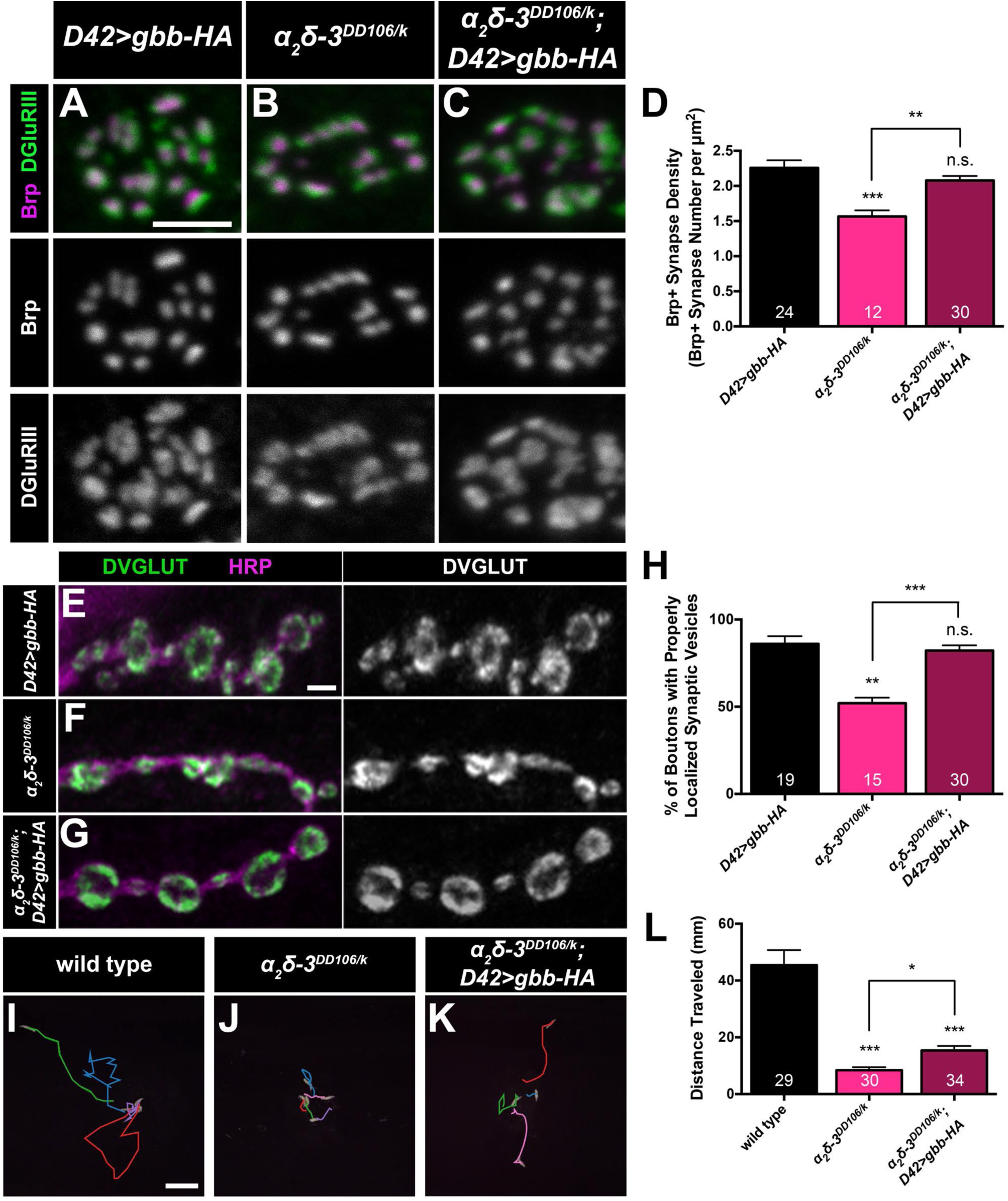
Gbb overexpression in motor neurons suppresses *α_2_δ-3* mutant phenotypes. (A-C) Representative z-projections of boutons of the indicated genotypes labeled with Brp (magenta) and DGluRIII (green). Scale bar: 2 µm. (D) Quantification of the Brp+ synapse density. *D42>gbb-HA*: 2.3 ± 0.1, *α_2_δ-3^DD106/k10814^*: 1.6 ± 0.1, *α_2_δ-3^DD106/k10814^; D42>gbb-HA*: 2.1 ± 0.1. (E-G) Representative z-projections of boutons of the indicated genotypes labeled with DVGLUT (green) and HRP (magenta). Scale bar: 2 µm. (H) Quantification of the percent of boutons exhibiting properly localized synaptic vesicles. *D42>gbb-HA*: 86.0 ± 4.4, *α_2_δ-3^DD106/k10814^*: 52.1 ± 3.2, *α_2_δ-3^DD106/k10814^; D42>gbb-HA*: 82.2 ± 3.0. (I-K) Representative traces of third-instar larvae of the indicated genotypes crawling for 3 minutes. Each color represents an individual larva. Scale bar: 10 mm. (L) Quantification of the total distance traveled by larvae of the indicated genotypes. n is the number of larvae scored. Wild type (*OregonR*): 45.5 ± 5.3, *α_2_δ-3^DD106/k10814^*: 8.4 ± 1.1, *α_2_δ-3^DD106/k10814^; D42>gbb-HA*: 15.4 ± 1.6. Error bars are mean ± SEM. n.s., not significantly different. *, p<0.05; **, p<0.01; ***, p<0.001.

Can motor neuron overexpression of Gbb also improve motor function of *α_2_δ-3* mutant animals? To address this question, we compared larval locomotion of *α_2_δ-3* mutants and *α_2_δ-3* mutants with neuronal Gbb overexpression. We find that *α_2_δ−3^DD106/k^* mutants exhibit dramatically reduced larval locomotion relative to wild-type larvae, moving only 18% of the distance as wild-type larvae over a three-minute interval **(Figure 8 I-J, L)**. Motor neuron-specific overexpression of Gbb results in a two-fold increase in the distance traveled by *α_2_δ−3^DD106/k^* animals **(Figure 8 I, K, L)**, consistent with improved synapse morphology in these animals. Thus, elevated levels of presynaptic BMP significantly improve larval locomotion in *α_2_δ-3* mutants. Given the separate requirement for α_2_δ-3 in calcium channel localization, it is striking that elevated autocrine BMP signaling improves larval locomotion of *α_2_δ-3* animals. Together, these data support the hypothesis that α_2_δ-3 facilitates BMP signaling.

### α_2_δ-3 limits diffusion of Gbb following its activity-dependent release

Extracellular proteins serve diverse functions in BMP pathways. They can antagonize BMP signaling by sequestering BMP ligands from their receptors, or alternatively, promote BMP signaling by serving as scaffolds to facilitate formation of ligand-receptor complexes (Sedlmeier and Sleeman, 2017). We hypothesized that the large glycosylated α_2_ peptide is well-positioned to modulate the extracellular distribution of Gbb following its activity-dependent release. In this case, we might observe a shift in extracellular Gbb in the presence and absence of α_2_δ-3.

To test this prediction, we utilized an assay of activity-dependent Gbb release (James et al., 2014). Presynaptic Gbb release is triggered by high K^+^ stimulation and quantified by comparing Gbb-HA intensity within the presynaptic membrane in unstimulated and stimulated preparations. Following stimulation, we find a roughly 50% decrease in mean Gbb-HA intensity within the presynaptic membrane, indicative of Gbb release **(Figure 9 A-B, G;** James et al., 2014). We next investigated if loss of α_2_δ-3 interferes with presynaptic release of Gbb in *α_2_δ-3* mutant backgrounds. We do not find alterations in presynaptic Gbb-HA levels in the absence of stimulation in either α_2_δ-3 mutant background. Moreover, stimulation leads to comparable reductions of intracellular Gbb-HA in both backgrounds **(Figure 9 A-G)**. The finding that presynaptic Gbb levels and activity-dependent release are independent of α_2_δ-3 is consistent with the model that α_2_δ-3 is an extracellular modulator of autocrine BMP signaling.

**Figure 9.**
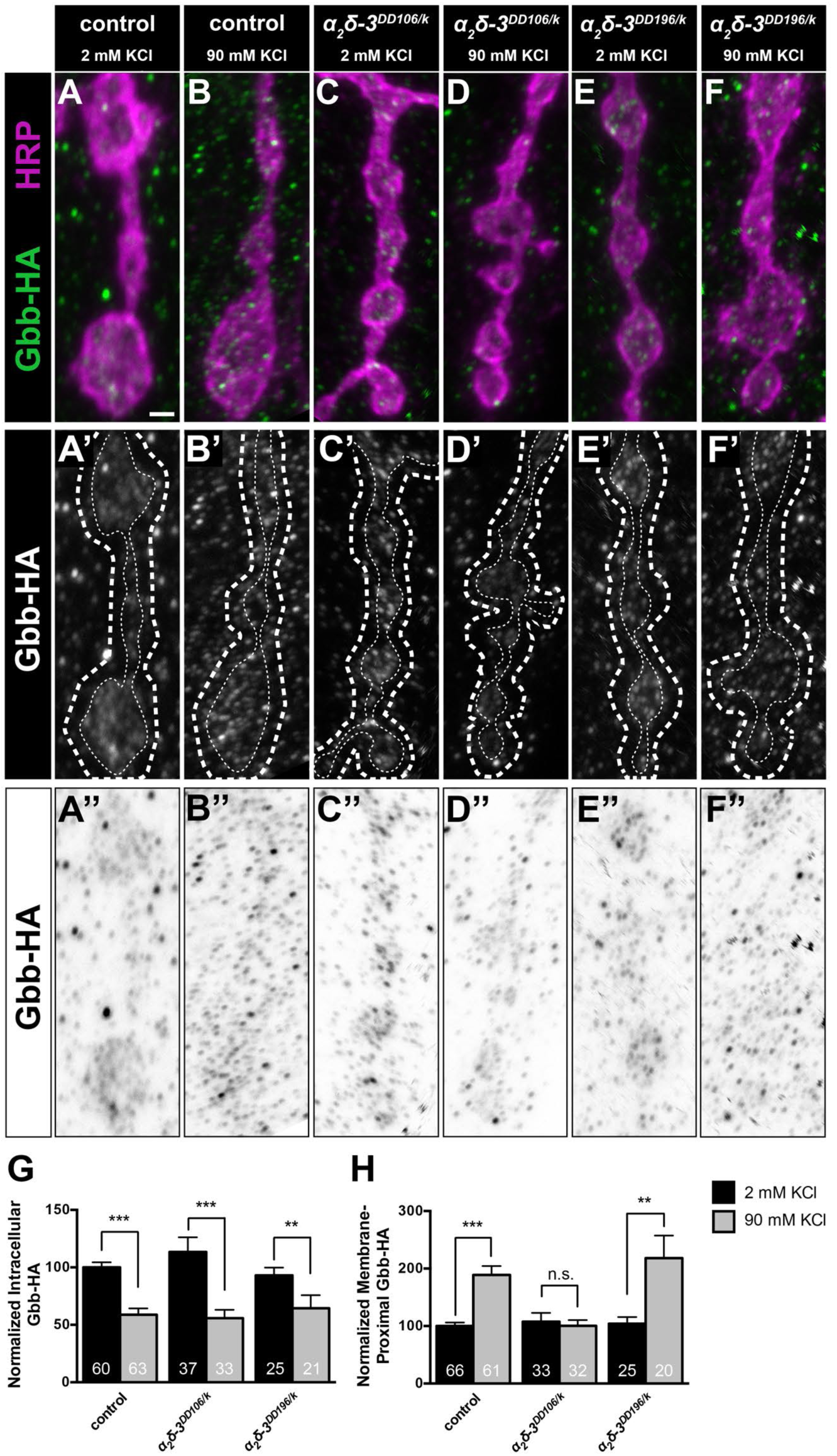
α_2_δ-3 limits extracellular diffusion of Gbb following its activity-dependent release. (A-F) Representative z-projections of boutons of the indicated genotypes labeled with HRP (magenta) and HA (green). Genotypes specifically are: control *(D42>gbb-HA)*, *α_2_δ-3^DD106/k^ (α_2_δ-3^DD106/k10814^; D42>gbb-HA)*, and *α_2_δ-3^DD196/k^ (α_2_δ-3^DD196/k10814^; D42>gbb-HA)*. Scale bar: 1 µm. (A’-F’) Individual Gbb-HA channel shown in grayscale. Membrane-proximal Gbb-HA was measured between the two dashed white lines. (A”-F”) Individual Gbb-HA channel with inverted colors. (G) Quantification of intracellular Gbb-HA normalized to control levels before (2 mM KCl) and after (90 mM KCl) neuronal stimulation. For all Gbb-HA release experiments, n is the number of NMJs scored. Control (2 mM): 100.0 ± 4.4, control (90 mM): 58.8 ± 5.4, *α_2_δ-3^DD106/k10814^* (2 mM): 113.5 ± 12.8, *α_2_δ-3^DD106/k10814^* (90 mM): 55.8 ± 7.2, *α_2_δ-3^DD196/k10814^* (2 mM): 93.0 ± 6.8, *α_2_δ-3^DD196/k10814^* (90 mM): 64.5 ± 11.4. (I) Quantification of membrane-proximal Gbb-HA normalized to control levels before and after neuronal stimulation. Control (2 mM): 100.0 ± 6.1, control (90 mM): 188.9 ± 15.6, *α_2_δ-3^DD106/k10814^* (2 mM): 107.6 ± 15.2, *α_2_δ-3^DD106/k10814^* (90 mM): 100.5 ± 9.8, *α_2_δ-3^DD196/k10814^* (2 mM): 104.1 ± 11.5, *α_2_δ-3^DD196/k10814^* (90 mM): 218.2 ± 39.2. Error bars are mean ± SEM. n.s., not significantly different. **, p<0.01; ***, p<0.001.

We then assessed if α_2_δ-3 acts subsequently to shape the distribution of Gbb following its activity-dependent release. At control NMJs, released Gbb-HA is visible as a cloud surrounding the presynaptic membrane following stimulation **(Figure 9 A-B”)**. We quantified extracellular Gbb by defining a 400-nm margin from the intracellular face of the presynaptic membrane as labeled by HRP into the extracellular space (between dashed lines in A’-D’). At control NMJs, we find a two-fold increase in mean Gbb-HA intensity in this membrane-proximal domain following stimulation **(Figure 9 H)** (James et al., 2014). In contrast, we do not detect an increase in membrane-proximal Gbb following stimulation at *α_2_δ-3^DD106/k^* mutant NMJs **(Figure 9 C-D”, H)**. To address a specific function for the extracellular α_2_ peptide, we analyzed the distribution of Gbb-HA following stimulation in *α_2_δ-3^DD196/k^* mutants since these mutants are predicted to retain the α_2_ peptide. Unlike *α_2_δ-3^DD106/k^* mutants, the extracellular distribution of Gbb-HA following stimulation is normal at *α_2_δ-3^DD196/k^* mutant NMJs **(Figure 9 E-F”, H)**. These findings argue that α_2_δ-3 serves as a physical barrier to Gbb diffusion to promote activity-dependent autocrine BMP signaling and further support a specific function for the extracellular *α_2_* peptide in the pathway.

## Discussion

In this study we uncovered a novel functional interaction between α_2_δ-3 and an activity-dependent, autocrine BMP signaling pathway at the Drosophila NMJ **(Figure 10)**. We find that α_2_δ-3 mutants display synaptic defects analogous to autocrine BMP mutants, and that α_2_δ-3 mutant phenotypes are suppressed by over-activating autocrine BMP signaling. We provide evidence that the α_2_ extracellular domain of α_2_δ-3 acts as a physical barrier to corral Gbb at the presynaptic membrane upon its activity-dependent release. Here we examine the relationship among α_2_δ proteins, autocrine BMP signaling, and the synaptic cleft microenvironment in maintaining synapse structure and function.

**Figure 10.**
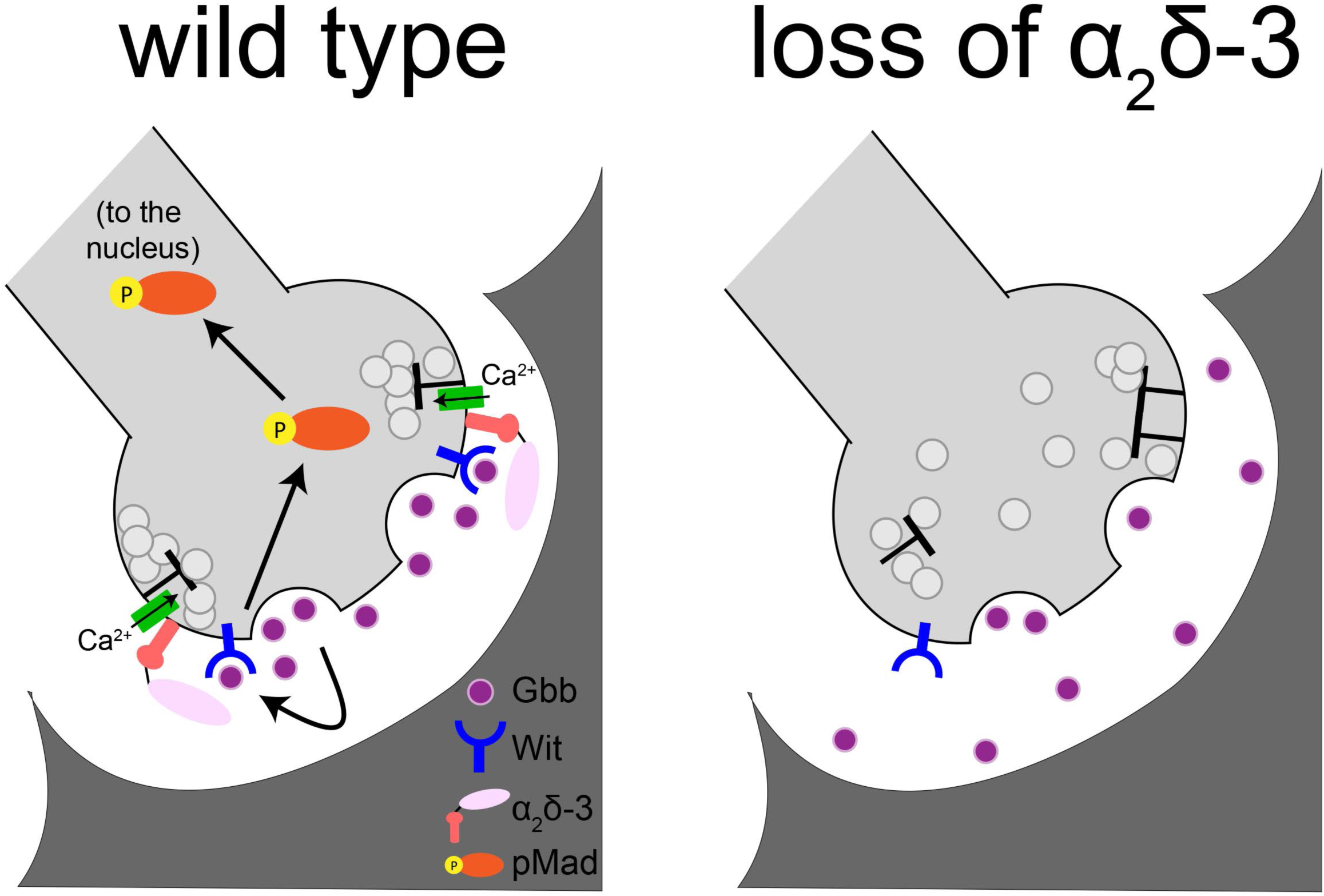
α_2_δ-3 promotes activity-dependent, autocrine BMP signaling. Proposed model for α_2_δ-3 acting as a physical barrier to promote Gbb signaling. Upon release from the neuron, Gbb is released into the synaptic cleft. With the aid of α_2_δ-3, Gbb remains in close proximity to the presynaptic membrane and is then able to activate the BMP Type II receptor Wit. The receptor complex in turn phosphorylates the transcription factor Mad, transducing a BMP signal back to the nucleus of the neuron.

### Multiple BMP signaling pathways at the Drosophila NMJ

BMP pathways regulate bouton number and morphology, bouton stabilization, presynaptic organization, neurotransmitter release, activity-dependent bouton addition, homeostatic plasticity, and postsynaptic differentiation at the Drosophila NMJ (this work)(Aberle et al., 2002; Ball et al., 2010; Eaton and Davis, 2005; Goold and Davis, 2007; James et al., 2014; Marques et al., 2002; McCabe et al., 2003; 2004; O’Connor-Giles et al., 2008; Piccioli and Littleton, 2014; Sulkowski et al., 2016). Multiple lines of evidence indicate that these pathways are at least partially separable. For instance, both bouton stabilization and activity-dependent bouton addition are non-canonical as they activate LIM kinase directly downstream of the Wit receptor (Eaton and Davis, 2005; Piccioli and Littleton, 2014). A second non-canonical BMP pathway regulates glutamate receptor composition at the NMJ. Interestingly, it requires local, presynaptic accumulation of pMad (Sulkowski et al., 2016). While accumulation of synaptic pMad depends on Wit, it is independent of Gbb, indicating that it is separable from the Gbb-dependent autocrine pathway described in this study.

How are the Mad-dependent pathways distinguished? In other words, how does retrograde BMP signaling regulate Mad-dependent gene transcription to promote scaling growth while autocrine BMP signaling regulates Mad-dependent gene transcription to maintain synapse structure and function? The distinct temporal requirement for the two pathways likely plays a key role to enable signal segregation. While the L1 stage is the critical period for the retrograde pro-growth signal, (Berke et al., 2013), we show that ongoing autocrine BMP signaling maintains synaptic structure. Thus, the signals are largely temporally separable. However, sustained muscle overexpression of Gbb during larval development fails to rescue baseline glutamate release indicating that temporal differences alone do not explain the distinct functions of the two pathways (Goold and Davis, 2007; McCabe et al., 2003). Moreover, we found that at L3, muscle-specific Gbb overexpression is less efficient at driving nuclear pMad localization than motor neuron Gbb overexpression **(Figure S3)**.

We propose that the temporal dynamics of activity-dependent release of BMP from presynaptic terminals promote robust induction of the downstream BMP cascade. This hypothesis is consistent with work demonstrating that pulsed TGF-β signaling dramatically increases the Smad transcriptional response (Sorre et al., 2014; Warmflash et al., 2012). This model is also supported by our work on the neuronal Gbb-binding protein Crimpy (Cmpy), which traffics Gbb to dense core vesicles (DCVs) for activity-dependent release (James et al., 2014). In the absence of Cmpy-mediated Gbb release, the ability of presynaptic Gbb to promote baseline neurotransmission is impaired. As expected, *cmpy* LOF mutants share some of the phenotypes described here including aberrant SSV distribution and disrupted embryonic bouton formation (Hoover and Broihier, unpublished). However, *cmpy* mutant *NMJs* are dramatically overgrown (James and Broihier, 2011), complicating analyses of synapse structure and function. Interestingly, Cmpy contains an IGF-binding domain and is poised to modulate the function of additional growth factors. Thus, here we focused on Gbb. Together, our findings provide a foundation for delineating how neurons discriminate among diverse secreted cues in their environment.

### BMPs, α_2_δ proteins, and the synaptic cleft microenvironment

The signaling range of BMP ligands in the extracellular space is tightly controlled by a large number of interacting proteins, including ECM components (Sedlmeier and Sleeman, 2017; Umulis et al., 2009; Wharton and Serpe, 2013). Not surprisingly, the ECM serves context-dependent functions in BMP signaling. For example, fibrillins sequester BMPs in the ECM to inhibit signaling (Sengle et al., 2008; 2011), while collagen type IV interacts with BMPs to augment local BMP signaling (Sawala et al., 2015; 2012; Wang et al., 2008). Specifically, collagen type IV immobilizes the BMP ligand Dpp to enable short-range signaling in Drosophila. More broadly, the ECM is proposed to generate a structural barrier through which BMP diffusion is limited (Sedlmeier and Sleeman, 2017). Such a barrier function could reasonably play a critical activating role in autocrine BMP pathways, such as the one described here.

Our findings argue that α_2_δ-3 likely serves as a diffusion barrier to Gbb following its activity-dependent release from neurons. α_2_δ proteins are present in the cleft and contain VWA domains, protein-protein interaction domains commonly found in extracellular scaffolds. The α_2_ subunit is glycosylated at multiple residues (Dolphin, 2012; 2018), raising the possibility that it could bind Gbb directly, since glycosylated proteins are well-established interactors of BMP ligands in the extracellular space that regulate their range and activity (Hayashi et al., 2009; Negreiros et al., 2018). Regardless of whether α_2_δ-3 and Gbb interact directly, or via an intermediary, we propose that α_2_δ-3 is a component of an extracellular scaffold that corrals BMP near the presynaptic membrane to facilitate the ligand-receptor interaction.

In mammals, α_2_δ-1 promotes excitatory synaptogenesis by acting as a postsynaptic receptor for glial thrombospondin (Risher et al., 2018; Eroglu et al., 2009). At the Drosophila NMJ, we found that overexpression of neuronal Gbb suppresses α_2_δ-3 mutant phenotypes **(Figure 8)**, indicating that α_2_δ-3 is not an obligate BMP receptor. Instead, we argue that α_2_δ-3 facilitates local BMP signaling. Is α_2_δ-3’s only function in the cleft local modulation of autocrine BMP signaling? Mammalian α_2_δ-1 binds a large number of synaptic proteins in addition to thrombospodin, including integrin subunits, neurexin, neuroligin, and cadherins (Brockhaus et al., 2018), suggesting that α_2_δ proteins interact widely with synaptic signaling or adhesion complexes. While we found clear phenotypic similarities between autocrine BMP mutants and α_2_δ-3 mutants, we could not analyze strong allelic combinations of α_2_δ-3 in late larval development as its function in Cac localization renders strong alleles late embryonic lethal. To determine if α_2_δ-3 modulates other extracellular signals at the Drosophila NMJ, it would be informative to generate α_2_δ-3 alleles that specifically delete the α_2_ domain to distinguish α_2_δ-3’s function in Cac localization from its Cac-independent roles in modulating extracellular signaling or adhesion.

### Autocrine signals can provide neurons with continuous feedback on their activity state

In this work, we demonstrate that an activity-dependent autocrine BMP pathway maintains synapse structure and function at the Drosophila NMJ. This pathway is spatially and temporally separable from the long-studied retrograde BMP pathway that initiates scaling growth (Berke et al., 2013; James et al., 2014; McCabe et al., 2003). Target-derived retrograde cues serve critical, conserved functions regulating neuronal survival, connectivity, and synaptogenesis (Zweifel et al., 2005). However, the broad expression patterns and pleiotropic phenotypes of many conserved signaling proteins has obscured a detailed view of their synaptic functions. The increasing ease of generating genetically mosaic animals, in which the protein-of-interest is deleted in a cell-type specific manner, has elucidated the cellular requirements of secreted synaptic signals, and revealed a number of autocrine pathways (reviewed in Herrmann and Broihier, 2018).

The autocrine BMP pathway we describe here is predicted to be transcriptional since it requires Mad. Thus, it does not appear to be a local synapse-specific signal, but rather a transcriptional cascade that regulates expression of genes involved in maintaining synapse structure and function. We propose that this activity-dependent autocrine cue provides ongoing feedback to motor neurons permitting them to maintain their activity state. It will therefore be essential to identify the cohort of Mad transcriptional targets induced by this activity-dependent cascade. Interestingly, BMP4 was recently found to be a presynaptic, autocrine cue in the mammalian hippocampus that signals through a canonical Smad pathway (Higashi et al., 2018). And similar to presynaptic Gbb in Drosophila, BMP4 is trafficked to DCVs and released in response to activity. However, in contrast to the synaptic maintenance function we describe here, BMP4 signaling destabilizes synapses for synaptic elimination. Thus, activity-dependent autocrine BMP signaling plays a conserved role in regulating synapse density during development, though whether the pathway has a stabilizing or destabilizing effect on synapses is context-dependent. It will be essential to determine the molecular mechanisms through which activity-dependent BMP signal transduction exerts its broad range of effects on synapse structure and function.

## Materials and Methods

### Fly stocks

The following stocks were used: *OregonR* (wild type), *BG57Gal4* (Budnik et al., 1996), *D42Gal4* (Parkes et al., 1998), *Elav-GeneSwitch* (Osterwalder et al., 2001), *gbb^1^* and *UAS-gbb9.1* (Khalsa et al., 1998), *gbb^2^* (Wharton et al., 1999), *UAS-gbb-RNAi* (Ballard et al., 2010), *wit^A12^*, *wit^B11^*, and *UAS-wit* (2002), *mad^12^* (Sekelsky et al., 1995), *mad^1^* (Takaesu et al., 2005), *cac^sfGFP-N^* (Gratz et al., 2019), *α_2_δ-*3^2^ (Ly et al., 2008), *α_2_δ-*3^DD106^, α_2_δ*-3^DD196^*, and *α_2_δ-3^k10814^* (Dickman et al., 2008), *UAS-gbb-1xHA* (James et al., 2014). All experiments were performed at 25°C.

### Immunohistochemistry and immunofluorescence

Wandering third-instar (L3) larvae were dissected and processed for immunofluorescence as previously described (James and Broihier, 2011). Briefly, larvae were dissected in PBS and fixed in either Bouin’s fixative for 5 min or 4% PFA in PTX (PBS and 0.1% Triton X-100) for 15 min (for Brp/GluRIII apposition and Gbb-HA experiments). Larval fillets were then washed in PTX, blocked in PBT (PBS, 1% BSA, and 0.1% Triton X-100), and incubated with primary antibody overnight at 4°C. For Gbb-HA experiments, samples were incubated with primary antibody for 2 hr at room temperature (RT) and overnight at 4°C. All samples were washed in PBT and incubated with secondary antibody for 2 hr at RT. The samples were then washed in PTX and mounted in either ProLong Diamond (Thermo Fisher Scientific) for Brp ring experiments or 60% glycerol for all other experiments.

For embryonic experiments, late-stage embryos were transferred using a small paintbrush to piece of double-sided tape. The embryos were rolled out of their chorion and placed on a plate in a small drop of PBS. Late-stage embryos have trachea fully formed and were removed from their vitelline membrane with forceps. Embryos were pinned at the head and tail using sharpened pins. Tungsten wire was used to pierce the body wall, and scissors were used to make a lateral cut down the stretched embryo. PBS was replaced with Bouin’s fixative for 5 min. The embryo was then rinsed with PBS and placed into a mesh basket surrounded by PBS in a small Petri dish. The embryos in the basket were washed three times for 15 min in PBT while slowly shaking. Embryos were blocked with PBT with 10% NGS (PBTN) for 10-30 min and incubated with primary antibody diluted in PBTN overnight at 4°C. Samples were then washed in PBT for 45 min while shaking, blocked in PBTN for 10-30 min, and incubated with secondary antibody diluted in PBTN for 2 hr at RT. Embryos were washed in PBT three times for 15 min each and PBS twice for 5 min each prior to being mounted in 60% glycerol.

For larval CNS experiments, wandering third-instar larvae were dissected and processed for immunofluorescence as previously described (Politano et al., 2019). Briefly, larvae were dissected in Schneider’s media (Sigma) supplemented with 10% FBS (Sigma) and 10U/mL Penicillin-Streptomycin (Thermo Fisher Scientific), then fixed in 4% PFA for 20 min. They were transferred to borosilicate tubes and washed in PBS and blocked in PTN (PBS, 0.1% Triton X-100, and 1% NGS) at RT on a rocker for at least 1 hr prior to overnight incubation in primary antibodies diluted in PTN at 4°C. Secondary antibody incubation in PTN was done for 3 hr at RT after primary antibody was washed out with PBS. Samples were mounted in ProLong Gold Antifade with DAPI (Thermo Fisher Scientific) after washing in PBS.

The following primary antibodies and concentrations were used: rabbit anti-Bruchpilot (NC82, Developmental Studies Hybridoma Bank) at 1:100, rabbit anti-GluRIII (gift from A. DiAntonio) at 1:2500, rabbit anti-DVGLUT (gift from A. DiAntonio) at 1:10000, rabbit anti-GFP (Abcam, ab6556) at 1:1000, rabbit anti-pMad (Cell Signaling, 41D10) at 1:100, guinea pig anti-Eve (gift from J. Skeath) at 1:2000, rat anti-HA (Roche, 11-867-423-001) at 1:200, and DyLight-594 anti-HRP (Jackson ImmunoResearch) at 1:300. The following goat species-specific secondary antibodies were used: Alexa Fluor 488, Alexa Fluor 568, and Alexa Fluor 647 (Invitrogen) at 1:300.

### RU486-GeneSwitch

Induction of Gal4-dependent transcription in GeneSwitch-Gal4 lines is induced by feeding animals with RU-486 since Gal4 is rendered steroid-inducible via fusion with the human progesterone receptor. Per Osterwalder et al., 2001, adults and their larval offspring were raised on non-RU486 food. L2 or early L3 larvae were transferred to molasses plates containing 25 mg/mL RU486 (mifepristone; Sigma Aldrich) and topped with a yeast paste made from 1 g dried yeast and 2 mL 50 mg/mL RU486 in water. RU486 was prepared as a stock solution at 10 mg/mL in ethanol. Control larvae were fed food containing equivalent concentrations of ethanol as vehicle at the same developmental stage. We ensured that in the absence of RU486, *gbb^−/−^; Elav-GS>Gbb* animals display active zone phenotypes similar to that observed in *gbb* nulls. Indeed, without drug, we saw identical structural phenotypes as observed in *gbb* nulls, confirming RU486-dependence of transgene expression.

### Gbb-HA release following neuronal stimulation

Each larva was pinned at the head and tail while in cold hemolymph-like HL3 saline. An incision was made laterally, and one pin was moved closer to the other, so the body wall can loosen. The HL3 was replaced with either cold 2 mM K^+^ or 90 mM K^+^ buffer for 5 min. The larva was next washed with cold PBS and quickly dissected.

### Image acquisition and quantification

Larval NMJs were imaged on a Zeiss LSM 800 confocal microscope with 100x 1.4 NA and 40x 1.3 NA oil-immersion objectives. Complete z-stacks were acquired with optimized settings to ensure that oversaturation did not occur. Confocal settings were held constant across all of the genotypes being tested in an experiment. After imaging samples for Brp rings, Zeiss ZEN deconvolution software was used on images. Maximum z-stack projections were created by ImageJ (National Institutes of Health). No modifications to any images were made prior to quantification. Larval NMJ4 boutons in segments A2-A4 were manually counted using an Axioplan 2 equipped with a Colibri 2 illumination system using the 100x 1.4 NA oil-immersion objective.

Synapses, defined as the tight apposition between Brp and GluRIII puncta, were manually scored at NMJ4 terminal boutons, and synapse density was calculated by the number of Brp+ synapses per bouton area in square microns. For all Brp^+^ synapse density quantifications, n is the number of NMJ4 boutons scored.

The diameters of isolated planar-oriented Brp rings were measured from confocal images of Brp-stained terminal boutons at NMJ4 that had been deconvolved using Zeiss ZEN software. A line was manually drawn through each Brp ring in ImageJ, and the Brp mean pixel intensity was plotted across the line. The distance from intensity maximum to intensity maximum was determined in the plot window, and this calculation was determined to be the Brp ring diameter (Owald et al., 2012). For all Brp ring analyses, n is the number of NMJ4 boutons.

DVGLUT localization was assessed by manually drawing a line through each bouton in ImageJ and DVGLUT mean pixel intensity was plotted across the line. If there were two peaks in the expression plot, and the trough of the plot was less than half of the smallest peak, the bouton was marked as having properly localized DVGLUT. We performed this analysis only on boutons larger than 2 microns in diameter, as smaller boutons were often completely filled with synaptic vesicles. The percent of boutons exhibiting properly localized synaptic vesicles compared to the total number of boutons analyzed at single NMJ4 was calculated. For all DVGLUT localization analyses, n is the number of NMJs scored. Concurrently, we measured the integrated pixel intensity over the same drawn line as an estimate of the number of synaptic vesicles. For all integrated DVGLUT analyses, n is the number of NMJ4 boutons larger than 2 microns in diameter.

To determine pMad intensity within motor neuron nuclei, VNCs were imaged at the same confocal settings and equal thickness maximum projections of the dorsal portion of the VNCs were made for each. Regions of interest were determined using Eve immunoreactivity. For each Eve+ motor neuron nucleus within a single VNC, the pMad and Eve pixel intensities were each measured. After subtracting background immunofluorescence from each channel, the pMad intensity was divided by the Eve intensity, providing a ratio of pMad to Eve intensity for each Eve+ motor neuron in a single VNC. Ratios were then averaged for each genotype. For the pMad analysis, n is the number of motor neuron cells scored.

To quantify intracellular Gbb-HA, mean pixel intensity was determined inside the inner neuronal membrane for samples incubated in either the 2 mM or 90 mM K^+^ buffer. Background HA fluorescence was averaged and subtracted from each projection. The neuronal membrane as defined by HRP staining is roughly 200 nm itself. To calculate Gbb-HA proximal to the neuronal membrane, Gbb-HA mean pixel intensity was measured between the intracellular face of the presynaptic membrane as labeled by HRP and a distance of 200 nm away from the extracellular face of the neuronal membrane, a distance of 400 nm in total. Again, background HA fluorescence was averaged and subtracted from each projection.

### Electron microscopy and analysis

Third-instar larvae were dissected and fixed for 2 hr in 2% glutaraldehyde and 2.5% paraformaldehyde. Following post-fixation in 1% osmium tetroxide, the specimen was dehydrated through a graded ethanol series and embedded in Eponate 12. The block was sectioned with a Leica EM UC6 ultramicrotome and stained with uranyl acetate and lead citrate. Images were collected on a FEI Tecnai G2 Spirit BioTWIN transmission electron microscope at the Cleveland Clinic. Electron micrographs of NMJs were taken from NMJ 6/7 and segments A2-A3 in at least two larvae of each genotype. Active zones were identified as linear electron densities found between pre- and postsynaptic membranes. High-magnification images (>125,000) were used to determine the presence of membrane ruffling and/or a T-bar, which was defined as an electron-dense rod surrounded by vesicles and localized to the presynaptic membrane.

### Electrophysiology

Neuromuscular junction sharp electrode recordings were performed as previously described (Bruckner, Gratz et al., 2012). Male third-instar larvae were dissected in 0.25 mM Ca^2+^ modified HL3 (70 mM NaCl, 5 mM KCl, 15 mM MgCl_2_, 10 mM NaHCO_3_, 115 mM sucrose, 5mM trehalose, 5mM HEPES, pH 7.2). Recordings were performed in 0.6 mM Ca^2+^ modified HL3 with on muscle 6 of abdominal segments A3 and A4 using sharp borosilicate electrodes filled with 3M KCl. Recordings were conducted on a Nikon FN1 microscope using a 40x 0.80 NA water-dipping objective and acquired using an Axoclamp 900A amplifier, Digidata 1550B acquisition system, and pClamp 11.0.3 software (Molecular Devices). Electrophysiological sweeps were digitized at 10 kHz and filtered at 1 kHz.

For each cell, mean miniature excitatory junctional potential (mEJPs) amplitudes were calculated using Mini Analysis (Synaptosoft) from all consecutive events during a one-minute recording in the absence of stimulation. EJPs were stimulated at a frequency of 0.2 Hz using an isolated pulse stimulator 2100 (A-M Systems), with intensity adjusted to consistently elicit compound responses from both type Ib and Is motor neurons. At least 25 consecutive EJPs were recorded for each cell and analyzed in pClamp to obtain mean amplitude. Quantal content was calculated for each recording using the ratio of mean EJP amplitude to mean mEJP amplitude. Recordings were limited to cells with a resting membrane potential between −50 mV and −85 mV and muscle input resistance greater than 5 MΩ.

### Locomotion assay

Prior to imaging, 150 mm Petri dishes were prepared, each with a thin layer of 1% agarose and black fountain pen ink (Pelikan) as a dye. L3 larvae were grown at 25°C and 4-7 larvae were then transferred to the middle of a Petri dish with a few drops of dH_2_O to keep the larvae moist. Larvae were imaged for 3 min. Movies were converted to 0.1 Hz (0.1 frames per second) and then larval crawling distance was measured every 10 seconds using the ImageJ plugin Manual Tracking. For the locomotion analysis, n is the total number of larvae scored per genotype in over four experimental trials.

### Statistical analyses

All statistical analyses were performed using Prism 6 (GraphPad Software). Each graph presents the data as mean ± SEM with the n value at the bottom of each column. All pairwise comparisons were made using a Mann-Whitney test. For determining significance between groups of three or more genotypes, a one-way ANOVA followed by Dunnett’s post hoc test was applied on normally distributed data. A nonparametric Kruskal-Wallis test followed by a Dunn’s multiple comparison test was performed on datasets that did not have a normal distribution. Analyses were conducted with the relevant Gal4 lines or heterozygotes as controls. In all figures, p-values are as follows: *p<0.05, **p<0.01, and ***p<0.001. No significant difference is denoted as n.s.

## Acknowledgements

We are indebted to Midori Hitomi at the Cleveland Clinic Imaging Core for her electron microscopy expertise and technical assistance. We thank Aaron DiAntonio, Thomas Schwarz, Jim Skeath, and Troy Littleton for fly strains and/or reagents. We also thank the Developmental Studies Hybridoma Bank for antibodies and the Bloomington Drosophila Stock Center for fly stocks. We thank Colleen McLaughlin for thoughtful discussions and feedback. We are grateful for the helpful comments from Pola Philippidou and Dan Jindal on the manuscript. This work was supported by National Institutes of Health (NIH) grants F31NS101763 to K.M. Hoover, R01NS078179 to K.M. O’Connor-Giles and R01NS095895 to H.T. Broihier. This research also received support from the NIH grant T32GM008056 awarded to the Cell and Molecular Biology Training Program at Case Western Reserve University.

---

**Figure S1.**
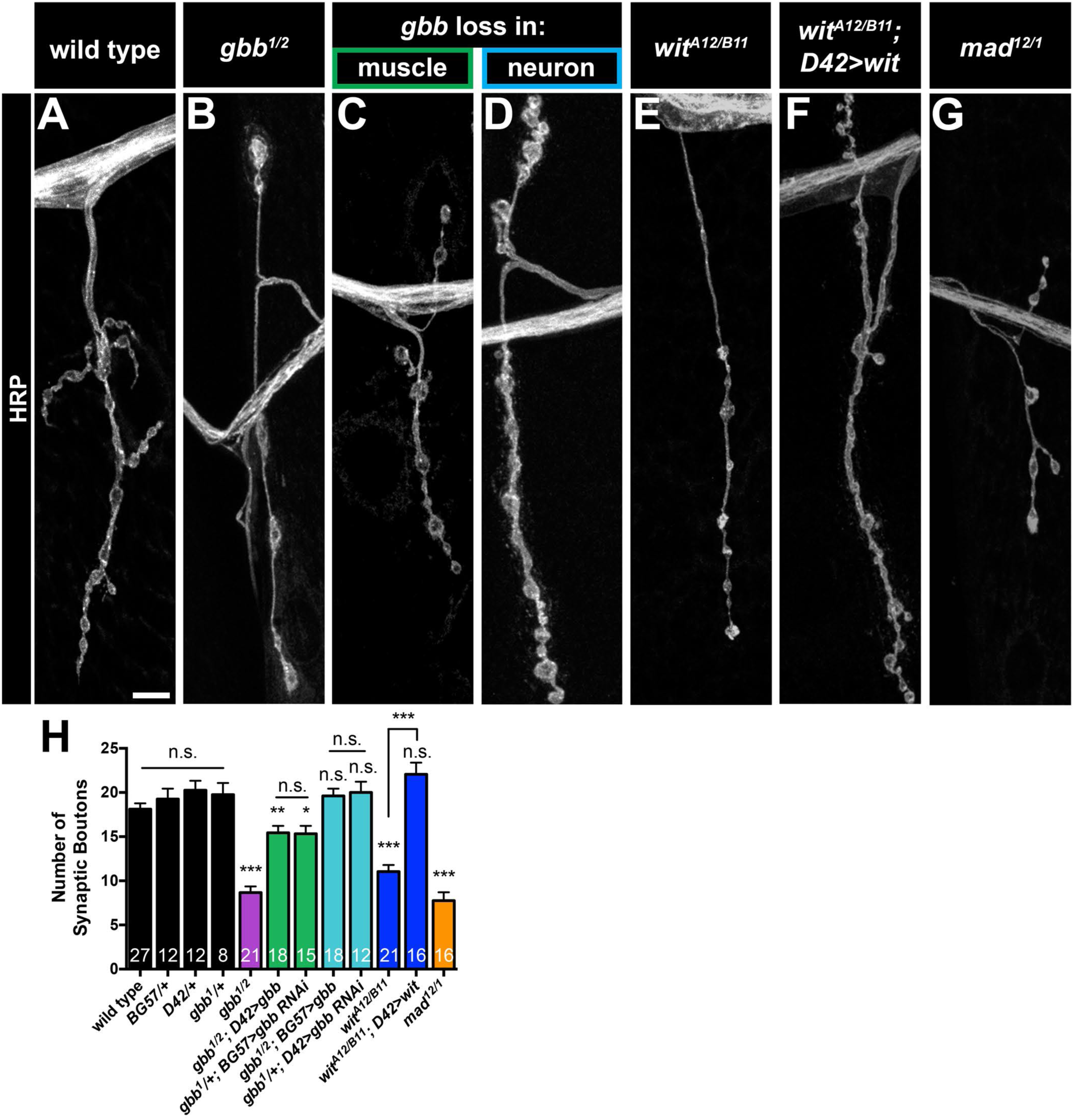
Presynaptic Gbb signaling does not regulate NMJ growth. (A-G) Representative z-projections of NMJs of the indicated genotypes labeled with HRP. Scale bar: 5 µm. (H) Quantification of the number of boutons. n is the number of NMJs scored. Wild type (*OregonR*): 18.1 ± 0.7, *BG57/+*: 19.3 ± 1.2, *D42/+*: 20.3 ± 1.1, *gbb^1^/+*: 19.8 ± 1.3, *gbb^1/2^*: 8.7 ± 0.7, *gbb^1/2^; D42>gbb*: 15.4 ± 0.8, *gbb^1^/+; BG57>gbb RNAi*: 15.3 ± 0.9, *gbb^1/2^; BG57>gbb*: 19.6 ± 0.8, *gbb^1^/+; D42>gbb RNAi*: 20.0 ± 1.2, *wit^A12/B11^*: 11.1 ± 0.7, *wit^A12/B11^; D42>wit*: 22.1 ± 1.3, *mad^12/1^*: 7.8 ± 0.9. Error bars are mean ± SEM. n.s., not significantly different. *, p<0.05; **, p<0.01; ***, p<0.001.

**Figure S2.**
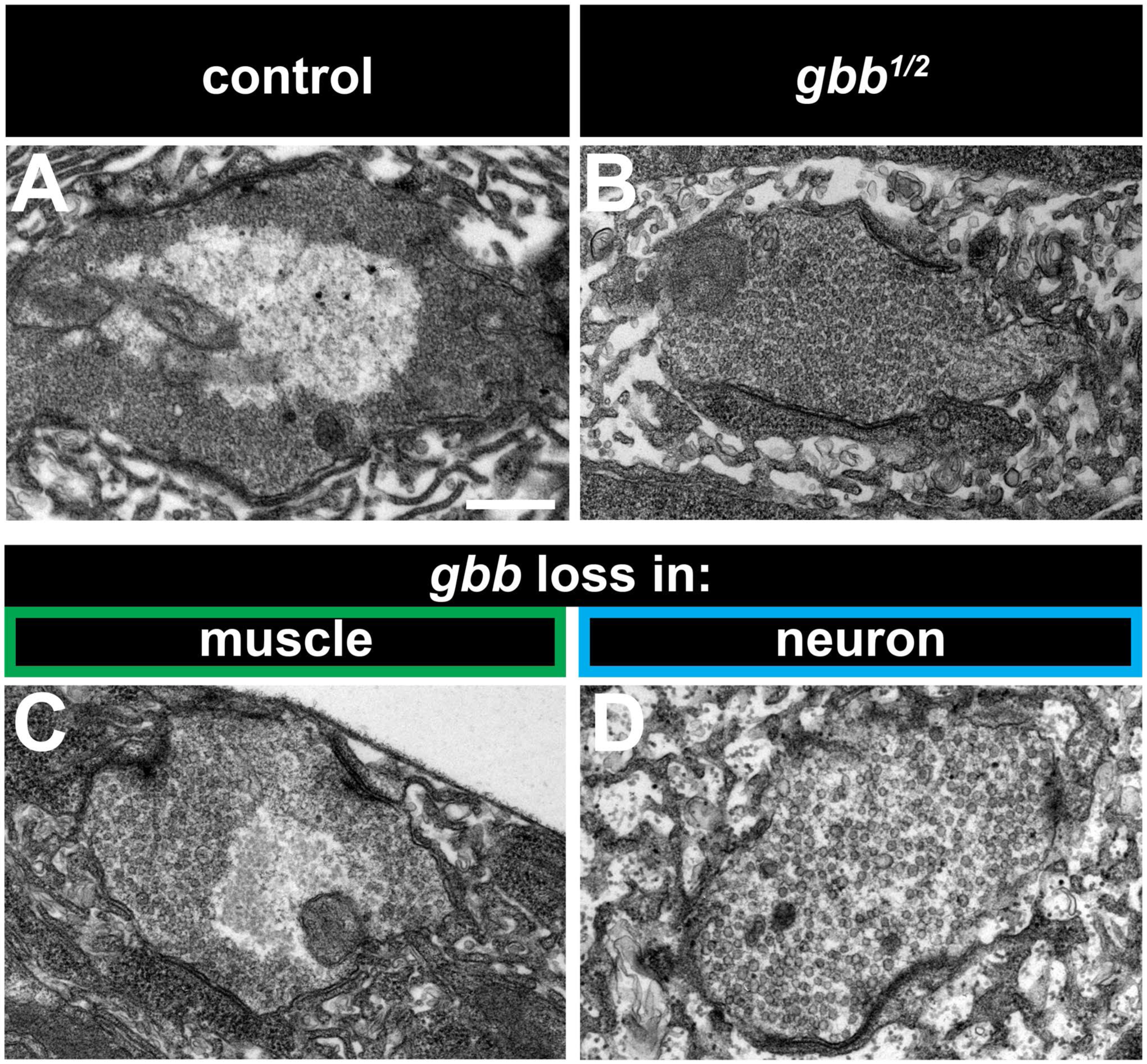
Presynaptic Gbb signaling regulates synaptic vesicle distribution within boutons at the ultrastructural level. (A-D) Representative transmission electron micrographs of boutons of the indicated genotypes. Scale bar: 1 µm. (A,C) Sections of control and muscle-specific *gbb* LOF boutons reveal synaptic vesicles localized to the bouton perimeter. (B,D) Sections of *gbb^1/2^* and neuron-specific *gbb* LOF boutons demonstrate synaptic vesicles filling the entire bouton.

**Figure S3.**
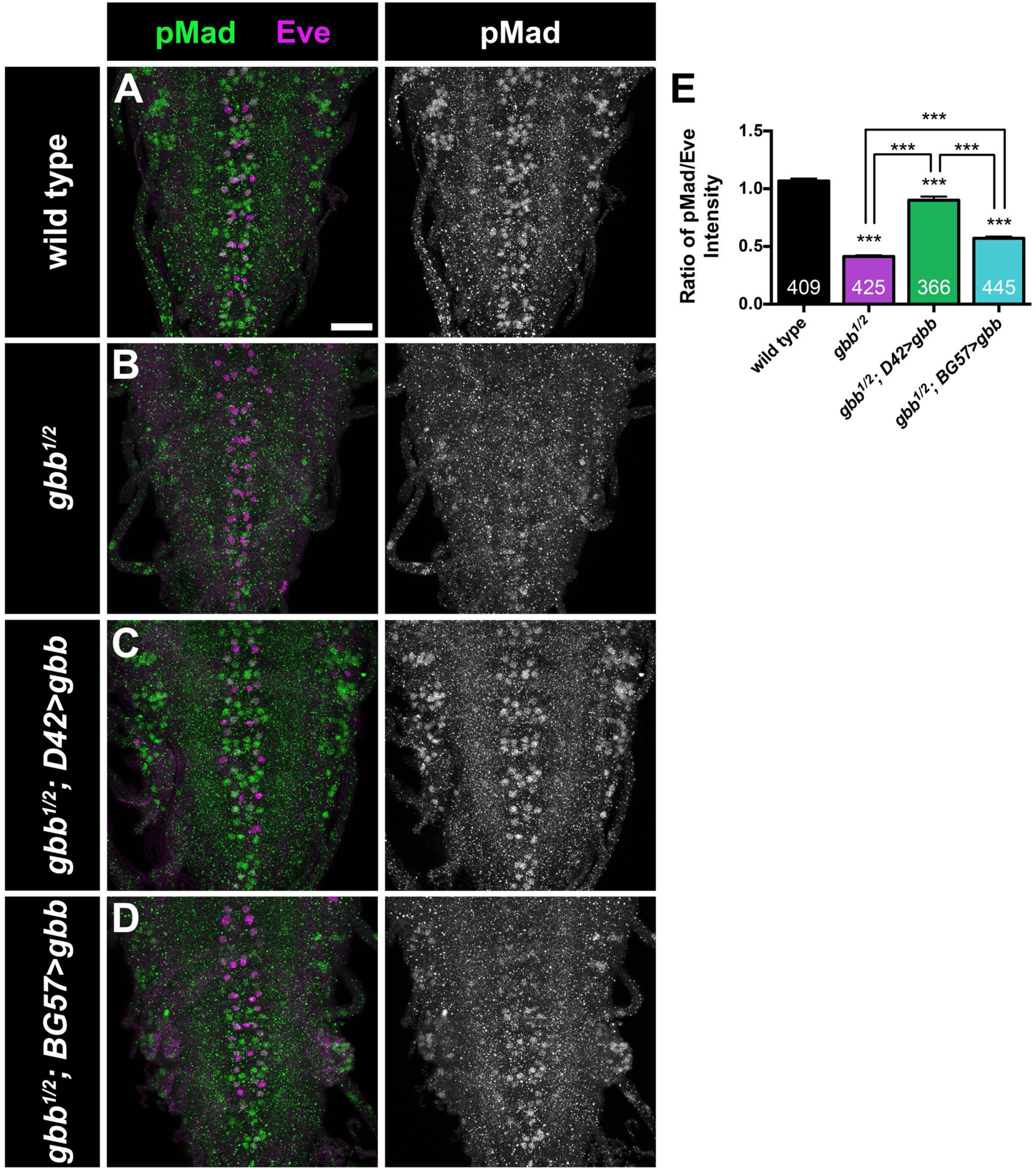
Presynaptic Gbb signaling is sufficient to promote pMad accumulation in the larval VNC. (A-D) Representative z-projections of VNCs labeled with Eve (magenta) and pMad (green) of the indicated genotypes. Scale bar: 25 µm. (E) Quantification of the ratio of pMad to Eve intensity in Eve+ motor neuron cells within the VNC. n is the number of motor neuron cells scored. Wild type (*OregonR*): 1.1 ± 0.0, *gbb^1/2^*: 0.4 ± 0.0, *gbb^1/2^; D42>gbb*: 0.9 ± 0.0, *gbb^1/2^; BG57>gbb*: 0.6 ± 0.0. Error bars are mean ± SEM. ***, p<0.001.

**Figure S4.**
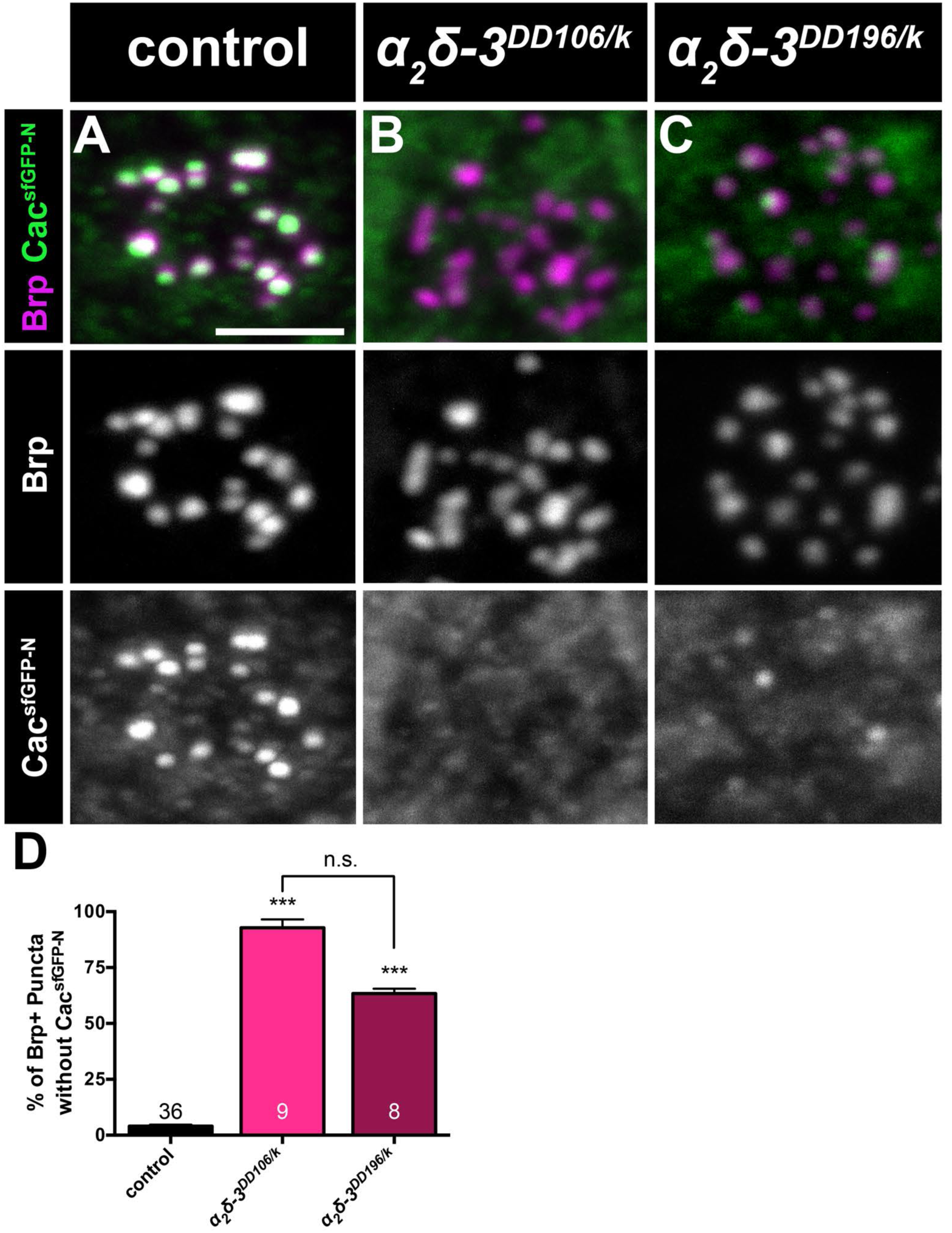
α_2_δ-3 is required for calcium channel localization to active zones. (A-C) Representative z-projections of boutons of the indicated genotypes labeled with Brp (magenta) and endogenously superfolder-GFP-tagged Cacophony (green). Scale bar: 2 µm. (D) Quantification of the percent of Brp+ puncta without Cac^sfGFP-N^. n is the number of boutons scored. Control (*cac^sfGFP-N^*): 4.0 ± 0.7, *α_2_δ-3^DD106/k10814^*: 92.8 ± 3.7, *α_2_δ-3^DD196/k10814^*: 63.4 ± 2.2. Error bars are mean ± SEM. n.s., not significantl

